# Astrocytic Slc4a4 regulates blood-brain barrier integrity in healthy and stroke brains via a NO-CCL2-CCR2 pathway

**DOI:** 10.1101/2023.04.03.535167

**Authors:** Qi Ye, Juyeon Jo, Chih-Yen Wang, Heavin Oh, Tiffany J. Choy, Kyoungin Kim, Angelo D’Alessandro, Yana K. Reshetnyak, Sung Yun Jung, Zheng Chen, Sean P. Marrelli, Hyun Kyoung Lee

## Abstract

Astrocytes play vital roles in blood-brain barrier (BBB) maintenance, yet how they support BBB integrity under normal or pathological conditions remains poorly defined. Recent evidence suggests pH homeostasis is a new cellular mechanism important for BBB integrity. In the current study, we investigated the function of an astrocyte-specific pH regulator, Slc4a4, in BBB maintenance and repair. We show that astrocytic Slc4a4 is required for normal astrocyte morphological complexity and BBB function. Multi-omics analyses identified increased astrocytic secretion of CCL2 coupled with dysregulated arginine-NO metabolism after Slc4a4 deletion. Using a model of ischemic stroke, we found that loss of Slc4a4 exacerbates BBB disruption and reactive gliosis, which were both rescued by pharmacological or genetic inhibition of the NO-CCL2 pathway *in vivo.* Together, our study identifies the astrocytic Slc4a4-NO-CCL2 axis as a pivotal mechanism controlling BBB integrity and repair, while providing insights for a novel therapeutic approach against BBB-related CNS disorders.

## Introduction

Disruption of blood-brain barrier (BBB) integrity is a devastating pathophysiological hallmark shared by various neurological disorders, often followed by severe consequences such as brain edema, neuronal damage, and eventually behavioral deficits (1, 2). Therefore, understanding the mechanisms underlying BBB integrity and developing strategies to reverse BBB damage will have far-reaching impacts on neurological diseases. Among the disorders accompanied by BBB dysfunction, ischemic stroke is a leading cause of death and disability worldwide (3). Although the primary aim of current stroke therapy is to restore cerebral blood flow, BBB dysfunction and associated secondary brain impairments remain difficult to treat (4) and the mechanisms underlying stroke-induced BBB disruption are still obscure.

While the core anatomic entities comprising the BBB are tightly connected endothelial cells, emerging evidence points to a crucial role of supporting cells in the neurovascular unit in BBB function (1, 5). Astrocytes are the most abundant and diverse glial cells in the central nervous system (CNS) (6, 7), and the interactions between astrocytic endfeet and blood vessel endothelial cells are essential for maintaining BBB integrity (5, 8). In particular, astrocytes secrete a number of paracrine factors that act on endothelial cells to regulate angiogenesis, blood flow, tight junction protein expression, and immune cell infiltration under normal physiological conditions (9). Among the paracrine factors secreted by astrocytes, chemokines and cytokines, via their downstream signaling pathways, play a critical role in the recruitment and migration of leukocytes into the brain during stroke and other disease conditions (10). There is now mounting evidence indicating that these signaling molecules may also directly modulate BBB permeability (11). While the unique anatomic feature and secretome-mediated inflammatory response of astrocytes are critical for BBB function, the regulatory mechanisms underlying astrocyte-BBB interaction in healthy and stroke brains remain poorly defined.

Among the growing list of pro-inflammatory chemokines involved in stroke, CCL2 (also known as MCP1) and its receptor CCR2 are particularly noteworthy because emerging clinical evidence suggests a positive correlation between the severity of stroke progression and CCL2 levels from patients’ serum and/or cerebrospinal fluid (12). However, the role of CCL2 in BBB damage after stroke remains obscure and the exact regulatory mechanism of CCL2 production is not defined. The surge of inflammatory factors after ischemic stroke is often accompanied by metabolic dysregulation(13). While a recent clinical study revealed a significant correlation between arginine metabolism dysregulation and CCL2 production in stroke patients (14), the molecular mechanism coupling the arginine metabolism pathway and CCL2 production is unknown.

To provide a favorable cellular environment, the extracellular and intracellular pH in the brain is strictly maintained by the balance of several ion species and the dynamic regulation of multiple ion transporters (15). Conversely, dysregulation of pH homeostasis in the brain is often implicated in a range of neurological disorders (16–18). CO_2_/HCO_3_^−^ is one of the most abundant and effective buffering ion pairs for pH control (15), and the imbalance of CO_2_/HCO_3_^−^ contributes to drastic pH reduction after ischemic stroke injury (19). While recent studies shed some light on the importance of astrocyte-specific acid extrusion in BBB recovery after stroke (20, 21), how astrocytes contribute to CO_2_/HCO_3_^−^ balance and related pH homeostasis in the context of BBB maintenance and repair are unknown. Previous literature has demonstrated that astrocyte-enriched electrogenic sodium-bicarbonate cotransporter 1 (Slc4a4) is responsible for shuttling HCO_3_^−^ across the cell membrane in a bidirectional manner to regulate both intra- and extracellular pH in response to intracellular signaling cascades as well as extracellular pH (22–25). In the clinical setting, Slc4a4 variants in patients are associated with a number of CNS disorders, including ischemic stroke (26–30). Despite its central role as an astrocytic pH regulator, whether and how Slc4a4 governs astrocyte-endothelial cell interaction in BBB maintenance and repair after stroke has not been investigated.

Here, we employed conditional genetic mouse models to investigate the role of Slc4a4 in both health and disease. Temporal ablation of Slc4a4 in astrocytes illustrated a requisite role of Slc4a4 in astrogenesis, astrocyte morphological complexity, and astrocytic Ca^2+^ physiology during normal development. In addition, loss of astrocytic Slc4a4 resulted in abnormal BBB structure and function in both normal and ischemic stroke brains. To identify mechanisms by which astrocytic Slc4a4 regulates BBB function, we performed unbiased multi-omics analyses of Slc4a4-deficient mice cortices and conditional media of Slc4a4-ablated astrocytes and discovered increased astrocytic CCL2 secretion upon loss of astrocytic Slc4a4. Moreover, our metabolomics profiling of Slc4a4-deficient cortices indicated that dysregulated arginine-nitric oxide (NO) metabolism is associated with increased CCL2 production. Further studies using pharmacological and genetic inhibition of either CCL2 or NO demonstrated a role of the Slc4a4-NO-CCL2 pathway in BBB recovery after stroke. Together, our study elucidates an indispensable role of Slc4a4 in astrocyte-BBB interaction, highlighting glial pH regulators as novel therapeutic targets for BBB-related pathology in ischemic stroke.

## Results

### Loss of astrocytic Slc4a4 reduces astrocyte morphological complexity in the brain

As a first step toward investigating the role of Slc4a4 in astrocytes, we assessed its expression in the mouse brain by performing fluorescence *in situ* hybridization and immunostaining. We found that Slc4a4 was expressed in the majority of astrocytes (Sox9+) in the cortex, but not neurons (NeuN+) (Figure 1A). Our analysis further revealed that >70% of Sox9-expressing astrocytes co-expressed Slc4a4 in the cortex during development and throughout adulthood (Figure 1B **and Supplemental Figure 1A**).

**Figure 1.**
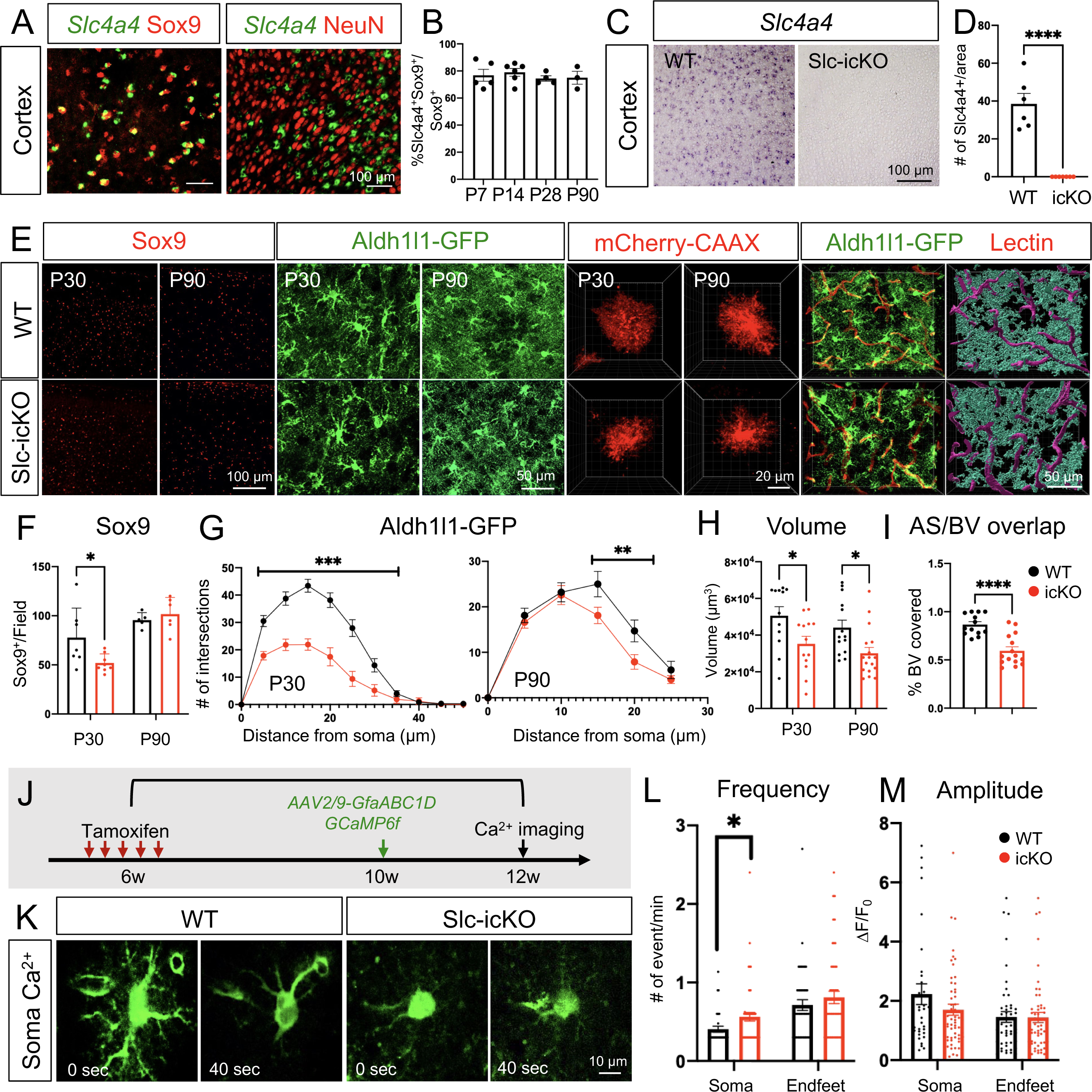
Slc4a4 is required for astrocytogenesis, morphological complexity and proper Ca^2+^ propagation. **(A)** Double *in situ*-immunofluorescence staining of Slc4a4 in astrocyte lineage (Sox9) and neuronal lineage (NeuN) in P28 mouse cortex. **(B)** Quantification of the number of Slc4a4-expressing astrocytes (Slc4a4+Sox9+) in cortex during development through adulthood. Data are presented as mean ± SEM. Each data point represents an individual animal. N = 3-5 for each age. **(C-D)** In situ hybridization confirms deletion of Slc4a4 in the cortex. **(E)** Immunofluorescence staining of astrocyte markers (Sox9, Aldh1l1-GFP) in the cortex from WT and Slc4a4-icKO mice at P30 or P90. Astrocyte morphology is labeled at single-cell resolution using AAV-PhP.eB -GfaABC_1_D-mCherry-CAAX by intracerebral injection at P1. Blood vessels were labeled by td-tomato lectin (red). Astrocyte-blood vessel interactions were reconstructed using IMARIS. **(F)** Quantification of the number of Sox9+ cells from WT and Slc4a4-icKO cortices at P30 and P90. Data are presented as mean ± SEM. Each dot indicates an individual animal. *p<0.05 by Student’s t-test. **(G)** Overall complexity of astrocytes (Aldh1l1-GFP) was measured by Sholl analysis. Data are presented as mean ± SEM. n = 6-8 cells per animal and N = 4-6 mice per genotype. **p<0.01, ***p<0.001 by two-way ANOVA. **(H)** Astrocyte volume was reconstructed and quantified using IMARIS software. Each dot represents an individual astrocyte. n = 14-18 astrocytes and N = 4 mice per group. **(I)** Quantification of blood vessel area covered by astrocytes after IMARIS 3D reconstruction. n = 3-4 images per animal and N = 3-4 mice per genotype. **p<0.01 by Student’s t-test. **(J)** Schematic of measuring astrocytic spontaneous calcium signaling in the cortex from WT and Slc4a4-icKO mice. Tamoxifen (100mg/kg) was injected for 5 consecutive days at 6 weeks old to induce gene deletion, followed by stereotaxic AAV2/9-GfaABC_1_D-GCaMP6f virus injection into the cortex at 10 weeks old. Astrocytic calcium from *ex vivo* cortical slices was imaged at 12 weeks old. **(K)** Representative images of astrocytic soma spontaneous calcium activity from WT and Slc4a4-icKO mice. **(L-M)** Quantification of amplitude and frequency of GCaMP6f signal events in cortical astrocyte soma and endfeet. Data are presented as mean ± SEM. Total number of cells is n = 30-40 collected from N = 6-8 mice of each genotype. Student’s t-test was used for statistics, *p<0.05.

To delineate how Slc4a4 functions in astrocyte development, we generated astrocyte-specific Slc4a4 conditional null mice by crossing Slc4a4-floxed (Slc4a4^F/F^) mice with an Aldh1l1-CreER driver line (31) (Aldh1l1-CreER; Slc4a4^F/F^, denoted as Slc4a4-icKO). We administered tamoxifen either during development (injection at P3 and harvested at P30) or in adulthood (injection at 6 weeks and harvested at 12 weeks) for analysis, confirming complete deletion of Slc4a4 in the cortex by in situ staining (Figure 1, C and D) and by real-time qPCR on FAC-sorted (Fluorescence Activated Cell) astrocytes (**Supplemental Figure 1B**). We first assessed the expression of established astrocyte markers. Relative to wild-type (WT) littermates, mice with Slc4a4 deletion during development, but not in adulthood, showed a 35% reduction in the number of Sox9+ astrocytes (Figure 1, E and F).

To visualize the morphological complexity of astrocytes, we further crossed Slc4a4-icKO mice with the Aldh1l1-EGFP reporter line (32) and performed high-resolution confocal imaging for GFP+ astrocytes. Sholl analysis (IMARIS software) revealed that Aldh1l1-GFP+ astrocytes from Slc4a4-icKO mice had markedly decreased branches compared with controls throughout brain development (up to 50% reduction) and adulthood (up to 28% reduction) (Figure 1, E and G). To assess the morphological changes of Slc4a4-depleted astrocytes in a more precise manner, we sparsely labeled cortical astrocytes with an adeno-associated virus (AAV) vector expressing membrane-targeted mCherry under the astrocyte-specific promoter GfaABC_1_D (AAV-PhP.eB-GfaABC_1_D-mCherry-CAAX) (33) in the mouse cortex at P1, followed by tamoxifen-induced Slc4a4 deletion during early development or adulthood. In both cases, we found that cortical astrocytes had a significantly smaller volume (28%) after Slc4a4 deletion compared to WT (Figure 1, E and H; **Supplemental Figure 1C**). Since the unique, complex, and bushy morphological features of astrocytes are required for their extensive contacts with endothelial cells (34), we measured the interaction between astrocytes and blood vessels using an Aldh1l1-GFP reporter and fluorescent lectin. We found that the reduction in branched morphology of cortical astrocytes in adult Slc4a4-icKO mice was coupled with a 32% decrease in the vasculature area covered by astrocytic endfeet processes, validated by 3D rendering (Figure 1, E and I). Together, these data indicate that Slc4a4 is required for astrocyte morphology and its association with blood vessels in the cortex.

### Slc4a4-depleted astrocytes exhibit abnormal Ca^2+^ wave propagation

Morphological changes noted in Slc4a4-deficient astrocytes raised the possibility that normal astrocyte physiology may be impaired. Previous studies have suggested a correlation between astrocytic morphological complexity and calcium (Ca^2+^) dynamics in the soma and processes, a direct indication of astrocyte physiological activity (21). To address whether astrocytic Ca^2+^ activity is altered in the absence of Slc4a4, we used an AAV vector to express a genetically encoded calcium indicator (AAV2/9-GfaABC_1_D-GCaMP6f) (35) in the cortex of adult Slc4a4-icKO and WT mice (Figure 1J). Two weeks after viral injection, we used established imaging algorithms to assess Ca^2+^ dynamics in the soma and endfeet of cortical astrocytes. While Ca^2+^ amplitudes in Slc4a4-deficient astrocytes remained unchanged in both soma and endfeet compared to WT astrocytes, we observed a slight increase (20%) of Ca^2+^ frequency in the soma, but not endfeet, of Slc4a4-deficient astrocytes (Figure 1, J-M; **Supplemental Figure 1D**). Together, these data indicate that loss of Slc4a4 moderately alters astrocyte Ca^2+^ physiology in the cortex.

### Loss of Slc4a4 results in hyper-vascularization coupled with junctional marker loss at the BBB

To investigate whether the observed morphological and physiological changes in Slc4a4-ablated astrocytes influence BBB function, we next evaluated whether the brain vasculature was altered in the absence of astrocytic Slc4a4. Using lectin to label blood vessels *in vivo*, we observed a 43% increase in blood vessel diameter and volume in Slc4a4-icKO mice compared to WT control (Figure 2, A-C), and a shift of blood vessel diameter distribution towards larger-sized vessels in Slc4a4-icKO mice (Figure 2D). Consistent with this enlarged blood vessel phenotype, we saw an up to 2-fold increase in endothelial cell marker expression (CD31 and Glut1) (Figure 2, A and E; **Supplemental Figure 2, A and B**). Notably, we observed a 50% upregulation of AQP4 in Slc4a4-icKO mice (Figure 2, A and E; **Supplemental Figure 2C**), a water channel known to be upregulated in response to osmotic imbalance by abnormal intracellular ion concentrations (e.g., Na^+^) (36). To investigate if enlarged cortical blood vessels in Slc4a4-icKO affect BBB function, we performed BBB leakage experiments *in vivo* by injecting molecules of different sizes (Figure 2I). Whereas no significant leakages of albumin (Evans blue) and 3kDa dextran were observed, we found a more than 3-fold increase of small molecules infiltrated into the brain parenchymal (MW = 0.5 kDa, EZ-Link Sulfo-NHS-Biotin) (Figure 2, J and K; **Supplemental Figure 2D**). Since biotins are known to diffuse into the brain parenchymal via the paracellular pathway upon endothelial tight junction disruption (37), we examined whether loss of astrocytic Slc4a4 affected the expression of endothelial tight junction markers. We observed a deficient coverage of blood vessels (CD31+) by either ZO-1 or Claudin-5 (35% and 15% decrease, respectively) (Figure 2, F-H) in Slc4a4-icKO cortices. Collectively, these data suggest a critical role of astrocytic Slc4a4 in regulating endothelial tight junctions and BBB paracellular transport in the adult brain.

**Figure 2.**
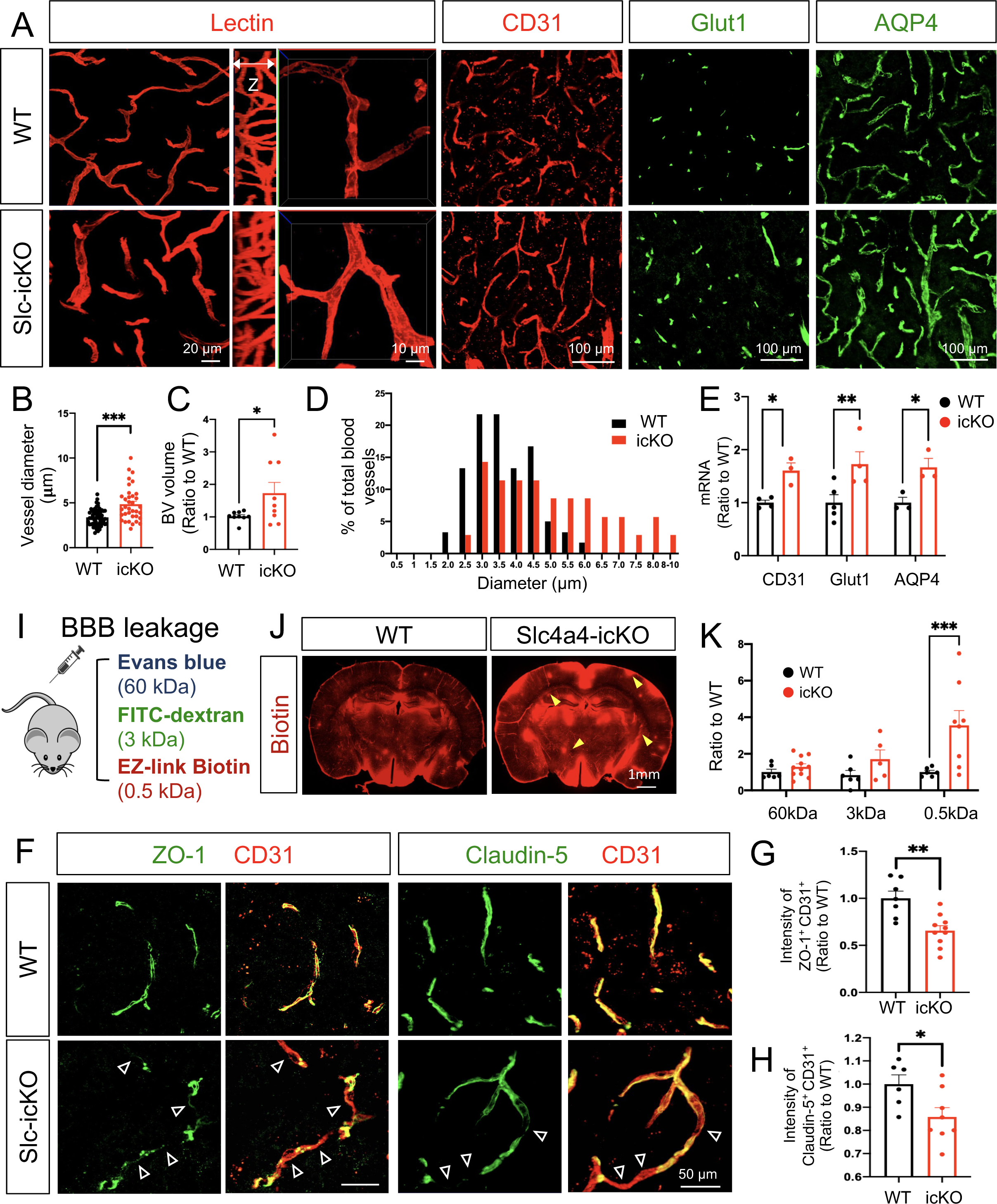
Loss of Slc4a4 results in hyper-vascularization coupled with junctional marker loss at BBB. **(A)** Blood vessel phenotype in Slc4a4-icKO cortex was examined by *in vivo* lectin labeling and immunofluorescence staining of vasculature markers (CD31, Glut1 and AQP4) at P90. **(B)** Quantification of blood vessel diameter by lectin labeling in WT and Slc4a4-icKO cortex. n = 35-60 vessels collected from N = 4-5 animals per genotype. ***p<0.001. **(C)** Quantification of blood vessel volume from IMARIS 3D reconstruction. Data are presented as mean ± SEM, n = 8-10 images from N = 4-5 animals per genotype. ***p<0.001 by Student’s t-test. **(D)** A histogram of average blood diameter measures in each genotype. n = 35-60 vessels collected from N = 4-5 animals per genotype. **(E)** qRT-PCR analysis of endothelial cell markers in the cortex from WT and Slc4a4-icKO mice. Data are presented as mean ± SEM. Each dot represents an individual animal (N = 3-5). *p<0.05 by Student’s t-test. **(I)** BBB leakage was assessed in WT and Slc4a4-icKO mice by intraperitoneal injection of Evans blue (indicates albumin leakage), intravenous injection of FITC conjugated dextran (3kDa), and intracardial injection of EZ-Link Sulfo-NHS-Biotin (indicates small molecule leakage) before harvest. **(J)** Representative images of stained biotin in the cortex from WT and Slc4a4-icKO mice. Yellow arrowheads indicate leakage of EZ-Link Sulfo-NHS-Biotin into the brain. **(K)** Extravasated Evans blue level in the brain was quantified by colorimetric assays. Extravasated FITC-dextran and EZ-Link Sulfo-NHS-Biotin were quantified based on intensity in brain sections. Data are presented as mean ± SEM. Each dot represents an individual animal (N = 6-8 per genotype). *p<0.05 by Student’s t-test. **(F)** Double immunofluorescence staining of tight junction markers (ZO-1, Claudin-5) and endothelial cell marker (CD31) in the cortex from WT and Slc4a4-icKO mice. Empty arrowheads indicate vessels missing coverage by tight junction proteins. **(G-H)** Quantification of the intensity of ZO-1 and Claudin-5 colocalized with CD31. Data are presented as mean ± SEM. N = 4-5 animals per genotype *p<0.05, **p<0.01 by Student’s t-test.

### Slc4a4-deficient astrocytes exhibit impaired BBB remodeling after ischemic stroke

To determine whether astrocytic Slc4a4 contributes to BBB remodeling after injury, we employed a cortical photothrombotic stroke (PTS) model (Figure 3A). We first assessed whether Slc4a4 is expressed in reactive astrocytes surrounding the lesion after various timepoints of cortical PTS. We found that Slc4a4 was highly expressed in S100b+ reactive astrocytes, suggesting its potential role after ischemic stroke (Figure 3B). We confirmed that Slc4a4 remains deleted in all the astrocytes surrounding lesion after stroke at 4 dpi (**Supplemental Figure 3A**). Next, we examined whether loss of Slc4a4 influences pH homeostasis after PTS using an extracellular pH-sensing probe pHLIP (pH-low insertion peptide) labeled with fluorescent dye ICG (pHLIP-ICG). It has been shown that the pHLIP probe can be washed out from the cell membrane in a pH-neutral environment (pH=7.2-7.4) but is inserted into the cell membrane when the extracellular environment turns acidic (pH=6-6.5) (38). As expected, IVIS imaging showed a strong signal as early as 1-day post-injury (1 dpi), but the signal was non-detectable in the non-injured brain (**Supplemental Figure 3B**), indicating an acidic extracellular milieu after stroke. The radius of acidic milieu was greater at the early stage after PTS (1 dpi vs. 4 dpi) (**Supplemental Figure 3B**). Furthermore, Slc4a4-icKO mice exhibited a 3-fold increase in extracellular acidity range compared to WT at 1 dpi (Figure 3C **and Supplemental Figure 3C**), suggesting that astrocytic Slc4a4 is required for maintaining pH homeostasis after stroke.

**Figure 3.**
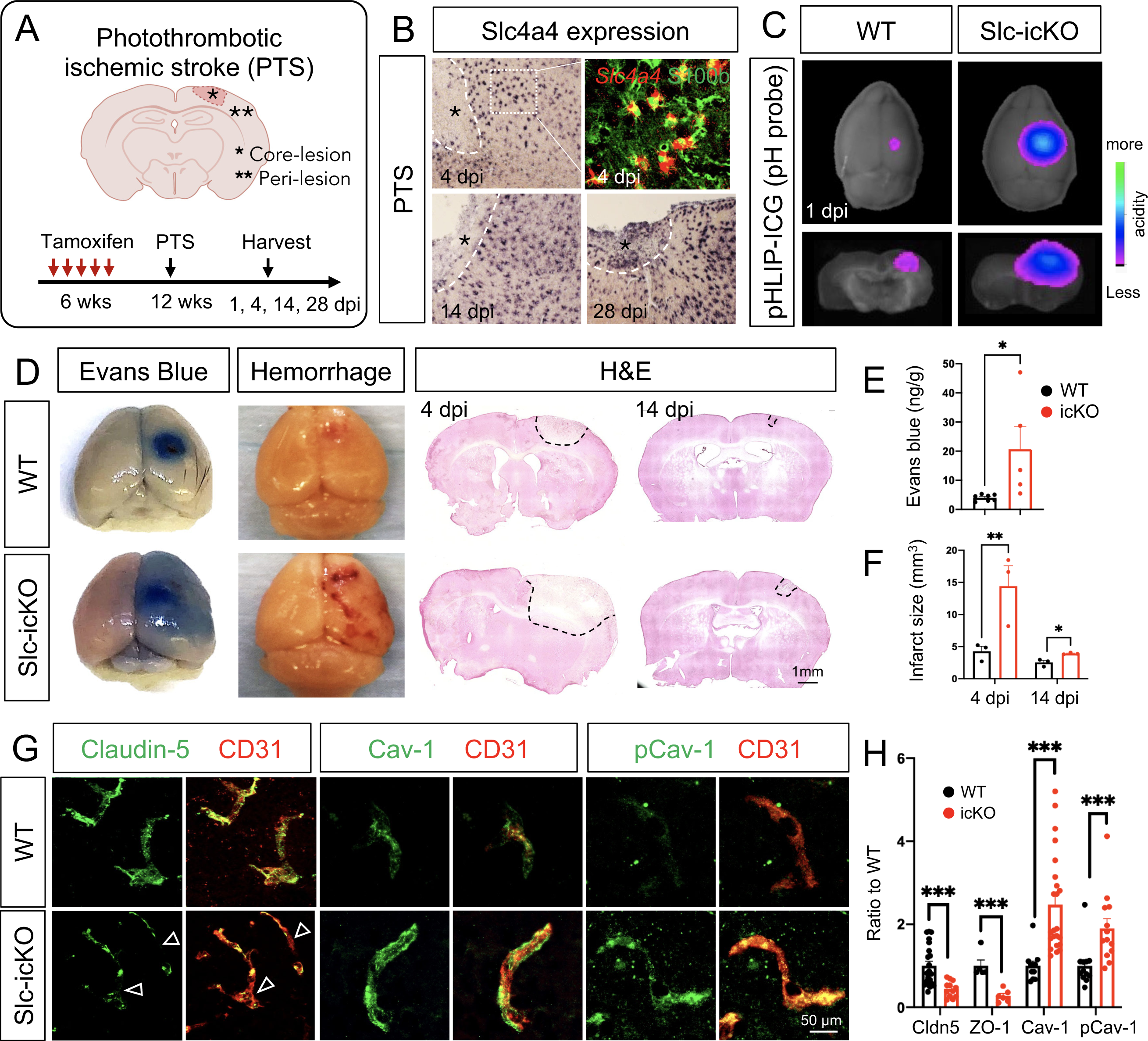
Slc4a4-deficient astrocytes exhibit impaired BBB remodeling after ischemic stroke. **(A)** Schematics of photothrombotic ischemic stroke (PTS) in WT and Slc4a4-icKO mice. Tamoxifen (100mg/kg) was injected for 5 consecutive days at 6 weeks old to induce gene deletion. PTS was induced at 12 weeks old (6 weeks post gene deletion), followed by brain harvesting at designated days post injury (dpi). Peri-lesion is defined as 150μm in distance from the lesion border. **(B)** Single or double staining of Slc4a4 (*in situ*) and S100b (immunostaining) in the WT cortex after PTS. **(C)** Extracellular pH in the stroke lesions was measured by intraperitoneal injection of pHLIP-ICG dye (1mg/kg). 24 hours after injection (1 dpi), brains from WT and Slc4a4-icKO mice were harvested and imaged using the Bruker Xtreme Imager with 735 nm excitation and 830 nm emission wavelength. **(D)** Representative images of albumin leakage (Evans blue), hemorrhage, and gross histology (H&E) of WT and Slc4a4-icKO mice at 4 or 14 dpi. **(E)** Evans blue levels in stroked brains from WT and Slc4a4-icKO mice were determined by colorimetric assays. Data are presented as mean ± SEM. Each dot represents an individual animal. N = 5-7 per genotype. *p<0.05 by Student’s t-test. **(F)** Quantification of infarct size is based on H&E staining of serial 40 μm-thick brain sections from stroked brains at 4 and 14 dpi. Each dot represents an individual animal. N = 3 per genotype. *p<0.05 by Student’s t-test **(G)** Double immunofluorescence staining of tight junction marker Claudin-5 and caveolae markers (Cav-1, pCav-1) with endothelial cell marker (CD31) at the peri-lesion area from WT and Slc4a4-icKO at 4 dpi. Empty arrowheads indicate loss of Claudin-5 (Cldn5). **(H)** Quantification of tight junctional markers (Cldn5, ZO-1) and caveolae markers (Cav-1, pCav-1) based on their intensity colocalized with CD31 in immunostaining. Data are presented as mean ± SEM. Each dot represents individual blood vessels collected from N = 4-6 animals per genotype. ***p<0.001, ****p<0.0001 by Student’s t-test.

We next examined whether exacerbated extracellular acidity in Slc4a4-icKO mice is associated with more severe pathological presentations after stroke. Strikingly, we observed a 5-fold increase in albumin leakage into the Slc4a4-icKO brains (Evans blue), along with an extensive hemorrhage area (Figure 3, D and E) and a 3-fold increase in infarct volume at 4 dpi (H&E) (Figure 3, D and F; **Supplemental Figure 3D**). The increased infarct volume in Slc4a4-icKO mice persisted until the stroke recovery stage at 14 dpi (Figure 3, D and F). The drastic BBB leakage at 4 dpi prompted us to examine the BBB structural integrity after stroke. We observed deletion of astrocytic Slc4a4 exacerbated tight junctional marker loss after stroke. In the Slc4a4-icKO brains, Claudin-5 expression was reduced by 56% at the peri-lesion area compared to WT controls, and unchanged in the core-lesion (Figure 3, G and H; **Supplemental Figure 4, A and C**). Another tight junction protein ZO-1 was also reduced by 75% in Slc4a4-icKO brains after stroke (Figure 3H **and Supplemental Figure 4B**). Since blood-borne proteins, such as albumin, cross the BBB via endothelial caveolae-mediated transcytosis, we further examined the expression of caveolin-1 (Cav-1), a structure protein of caveolae, and its phosphorylated form (pCav-1), an indicator of endothelial transcytosis activity (39). Corresponding to increased albumin leakage in the Slc4a4-icKO brains after stroke (Figure 3, D and E), we found a 2-fold increase in both endothelial Cav-1 and pCav-1 (Figure 3, G and H), suggesting elevated caveolae-mediated transcytosis. Taken together, our results demonstrate that the astrocytic Slc4a4 is required for re-establishing pH homeostasis and recovering BBB function after stroke.

### Loss of astrocytic Slc4a4 dampens reactive astrogliosis and astrocyte-endothelial interaction after stroke

Given the critical role of Slc4a4 in astrocytic interaction with endothelial cells and the key function of reactive astrocytes in BBB reconstruction after injury, we evaluated the role of Slc4a4 in reactive gliosis after stroke. Analysis of reactive astrocyte markers at 4 dpi showed a 40-50% reduction of GFAP expression in astrocytes surrounding the infarct lesion in the Slc4a4-icKO compared with WT mice after PTS (Figure 4, A and C). Further morphological analysis revealed a 25% reduction in total branch volumes of reactive astrocyte (GFAP+) in Slc4a4-icKO mice at 4 dpi (Figure 4, A and D). In addition, we observed a 30% decrease in the overall number of reactive astrocytes (S100b+) surrounding the lesion area of Slc4a4-icKO mice at 4 dpi (Figure 4, A and E). To label proliferating astrocytes, we administered multiple doses of BrdU (200mg/kg, every 12 hours from 1-3 dpi). Strikingly, we found an almost 100% reduction in proliferating astrocytes (S100b+; BrdU+) at peri-lesion areas of Slc4a4-icKO mice, indicating impaired local astrocyte proliferation after stroke. Since subventricular zone (SVZ) neural stem cells give rise to a subpopulation of reactive astrocytes in the stroke-lesioned area to participate in glial scar formation (40), we further investigated whether reduced SVZ astrocyte proliferation decreases the number of reactive astrocytes surrounding stroke lesions, and found an 80% reduction in proliferating SVZ astrocytes (BrdU+; Sox9+, Figure 4, A and G). At the chronic stage of stroke (14 dpi), we did not observe any proliferating astrocytes at the peri-lesion area nor the SVZ zone in either WT or Slc4a4-icKO mice (**Supplemental Figure 5A**). Nevertheless, we found that persistent dampened reactive gliosis, including a thinner glial scar, decreased S100b expression and reduced the number of astrocytes surrounding the lesion (**Supplemental Figure 5, A-D**). Together, these data indicate that loss of Slc4a4 impairs both local and SVZ astrocyte proliferation after cortical ischemia, which further leads to dampened reactive astrocyte response surrounding the lesion in both acute and chronic stages of ischemic stroke.

**Figure 4.**
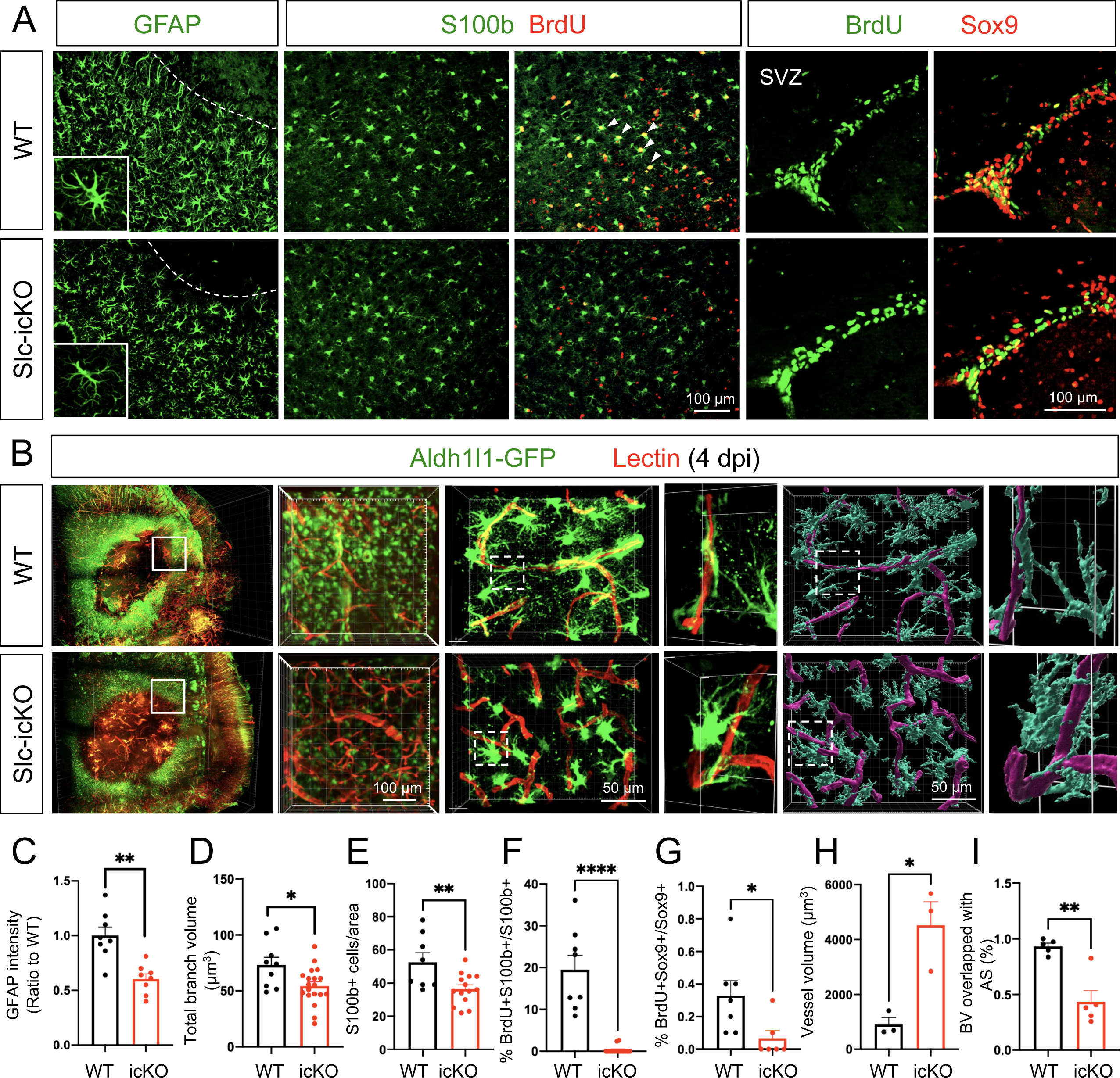
Loss of astrocytic Slc4a4 dampens reactive astrogliosis and astrocyte-blood vessel interaction after stroke. **(A)** Immunostaining of reactive astrocyte markers (GFAP, S100b) at the peri-lesion area at 4 dpi. To label proliferating astrocytes, WT and Slc4a4-icKO mice were intraperitoneally injected with BrdU (200mg/kg) every 12 hours from 1 to 3 dpi. S100b+ cells are co-labeled with BrdU to indicate local astrocyte proliferation. SVZ Sox9+ cells are co-labeled with BrdU to indicate SVZ astrocyte proliferation. **(B)** Whole-mount images of CLARITY-cleared mice brain at 4 dpi of PTS. Astrocytes were genetically labeled with Aldh1l1-GFP (green), and blood vessels were labeled with tomato-lectin (red). 40µm-thick sections were used for further IMARIS 3D reconstruction to visualize astrocyte-blood vessel interaction. **(C)** Quantification of GFAP intensity from immunostaining. Data are presented as mean ± SEM. Each dot represents an individual animal. N = 8 per genotype, **p<0.01 by Student’s t-test. **(D)** Quantification of total branch volume of reactive astrocytes from GFAP immunostaining. Data are presented as mean ± SEM. n = 9-19 cells collected from N = 3-5 mice per genotype. *p<0.05 by Student’s t-test. **(E)** Quantification of S100b+ cell number from immunostaining. Data are presented as mean ± SEM. Each dot represents an individual animal. N = 4-6 per genotype. **p<0.01 by Student’s t-test. **(F)** Quantification of proliferating reactive astrocytes at peri-lesion area (S100b+; Ki67+). Data are presented as mean ± SEM. Each dot represents an individual animal. N = 5-7 per genotype. ****p<0.0001 by Student’s t-test. **(G)** Quantification of the number of proliferating SVZ astrocytes (BrdU+; Sox9+). Data are presented as mean ± SEM. Each dot represents an individual animal. N = 4-7 per genotype. *p<0.05 by Student’s t-test. **(H-I)** Quantification of blood vessel volume and volume covered by astrocyte processes in the peri-lesion area from WT and Slc4a4-icKO cortices at 4 dpi. Each dot represents an individual animal. N = 3-4 per genotype. *p<0.05, **p<0.01 by Student’s t-test.

The foregoing data implicate that Slc4a4 modulates astrocyte morphology under both normal conditions and after ischemic stroke (Figure 1 **and** Figure 4A). To test whether loss of astrocytic Slc4a4 results in defective physical interaction between reactive astrocytes and blood vessels after stroke, fluorescent lectin-perfused Slc4a4-icKO and WT mice brains were subjected to CLARITY-based tissue clearing followed by light-sheet microscopy. Consistent with 2D histological staining in Figure 3D, 3D reconstruction of whole-mounted brains revealed larger infarct areas in Slc4a4-icKO mice with enlarged surrounding blood vessels (Figure 4, B and H). Of note, the blood vessel enlargement by loss of astrocytic Slc4a4 was much more pronounced after stroke compared to normal conditions (5-fold vs. 1.7-fold). Further confocal imaging of brain sections, followed by IMARIS 3D-rendering, showed a marked reduction of blood vessel area covered by astrocyte processes (53% decrease after stroke vs. 32% without stroke) (Figure 4, B and I). This observation suggests that astrocytic Slc4a4 is critical for astrocyte-blood vessel interaction after ischemic stroke.

### Slc4a4 regulates astrocyte-BBB integrity via astrocyte-derived cytokine CCL2

Emerging evidence showed a crucial role of astrocyte-derived components in angiogenesis and BBB function under both physiological and ischemic stroke injury conditions (41). To interrogate the mechanisms by which astrocytic Slc4a4 regulates astrocyte-BBB interactions, specifically astrocyte-endothelial interactions, we focused on astrocytic secretory factors. We first exposed a mouse endothelial cell line (bEnd3) to conditioned media (CM) from either WT (WT-CM) or Slc4a4-deficient astrocytes (KO-CM) (**Supplemental Figure 6A**). We observed that endothelial cell size (CD31) was enlarged by ∼60% and the number of proliferating endothelial cells (BrdU) was increased by 2-fold in the KO-CM treated group (**Supplemental Figure 6, B-D**). We then examined whether this *in vitro* system recapitulated the decreased junctional marker expression seen in Slc4a4-icKO mice. Indeed, we found a reduction of junctional marker ZO-1 expression upon treatment with KO-CM, phenocopying the loss of tight junction marker expression in Slc4a4-icKO mice *in vivo* (**Supplemental Figure 6, B and E**).

To directly examine whether astrocytes lacking Slc4a4 impact BBB permeability via astrocytic secretory factors, we performed co-culture assays and evaluated the transendothelial electrical resistance (TEER) across this bilayer, an established metric of paracellular BBB permeability *in vitro* (**Supplemental Figure 6F**). To this end, we co-cultured astrocytes with a monolayer of endothelial cells in the transwell, where astrocytes and endothelial cells were plated at the bottom of the well and in the insert respectively, exposing endothelial cells to astrocytic secretory factors without direct physical contact (**Supplemental Figure 6F**). The electro-resistance of the endothelial cell monolayer was measured as an indicator of their paracellular permeability. Interestingly, co-culture with Slc4a4-deficient astrocytes increased the permeability of the endothelial cells, as indicated by lower electric resistance (**Supplemental Figure 6G**). This observation demonstrates that Slc4a4 regulates astrocyte-mediated maintenance of BBB integrity via paracrine signaling factors, independent of direct physical interactions with endothelial cells.

To further elucidate the paracrine signaling molecules sensitive to Slc4a4 loss-of-function, we collected WT-CM and KO-CM for mass-spec profiling (LC-MS/MS) followed by cross-comparison with angiogenesis array analysis. Our analysis revealed a group of candidate proteins that function in angiogenesis and BBB function (Figure 5, A and B). Since Slc4a4-icKO mice brains showed an increased expression of CD31 (Figure 2A), which is often seen under inflammation conditions (42), we further analyzed the chemo/cytokine profile in astrocyte CM. Strikingly, we found an overall elevation of cytokines in KO-CM (Figure 5C). Among those cytokines, CCL2 was particularly interesting as it was also one of the top candidates from LC-MS/MS profiling (Figure 5B). To validate a potential astrocytic Slc4a4-CCL2 cascade, we quantitively measured CCL2 levels *in vitro* and *in vivo*. ELISA showed that CCL2 was enriched in conditioned media from Slc4a4-depleted astrocytes under both normal (50% increase) and oxygen-glucose deprivation (OGD) conditions (70% increase), the latter mimicking *in vivo* ischemic stroke (Figure 5D). Existing knowledge suggests that CCL2 promotes endothelial permeability by activating the endothelial CCR2 receptor (43); therefore we hypothesize that loss of Slc4a4 stimulates astrocytic CCL2 secretion and activates endothelial CCR2 receptor to induce BBB leakage (Figure 5E). To test this hypothesis, we measured cortical CCL2 levels, which were upregulated more than 2-fold in Slc4a4-icKO mice under normal and ischemic stroke conditions compared to WT (Figure 5F). Moreover, we confirmed increased astrocytic CCL2 expression by immunofluorescence co-labeling and found a more than 2-fold increase of CCL2 expression in the GFAP+ and S100b+ astrocyte populations at 4 dpi (Figure 5, G and I, **Supplemental Figure 7, A and B**). CCR2 was highly expressed in the endothelial population (CD31+) after stroke and was upregulated 6-fold at the peri-lesion area in Slc4a4-icKO mice brains at 4 dpi (Figure 5, H and I). In contrast, we did not observe astrocytic CCR2 expression at 4 dpi (Aldh1l1-GFP+; CCR2+, Figure 5I). Together, these data suggest that the Slc4a4-CCL2-CCR2 axis regulates endothelial function in a cell-nonautonomous manner.

**Figure 5.**
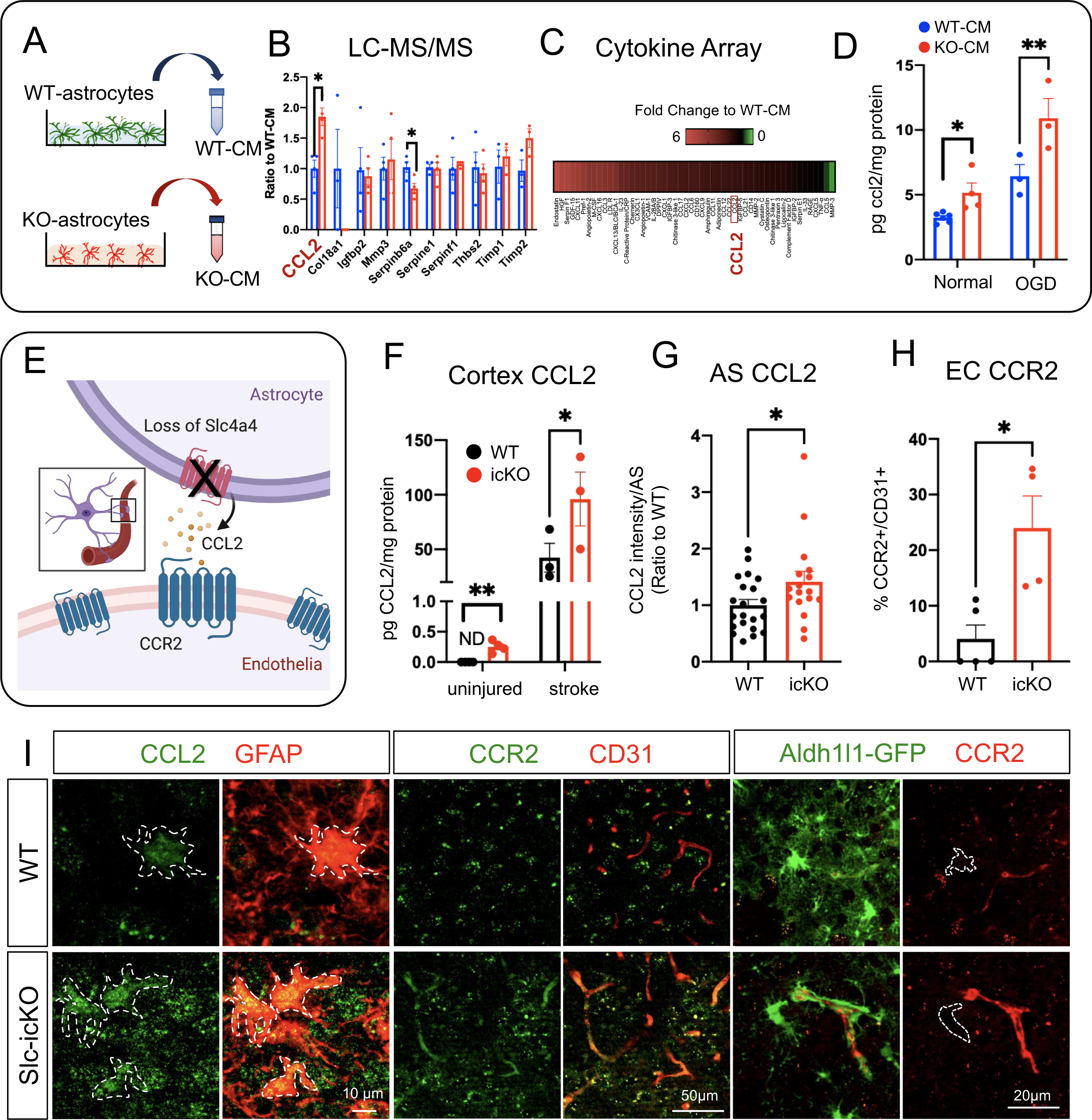
Loss of Slc4a4 upregulates astrocytic CCL2 and endothelial CCR2 after ischemic stroke. **(A)** Conditioned media (CM) was collected from primary WT and Slc4a4 KO astrocytes and subjected to LC-MS/MS-based unbiased proteomics and cytokine/chemokine array. **(B)** Angiogenic factors detected from LC-MS/MS-based unbiased proteomics. **(C)** Cytokine/chemokines that are changed in the CM from WT and Slc4a4 KO astrocytes. **(D)** CM from primary WT and Slc4a4 KO astrocytes cultured under either normal or oxygen-glucose deprivation (OGD) condition was collected for CCL2 measurement by ELISA. Each dot represents conditioned media collected from individual animals’ astrocytes. N = 3-5 per genotype. *p<0.05, ****p<0.01 by Student’s t-test. **(E)** Proposed model for the Slc4a4-CCL2-CCR2 axis regulating astrocyte-endothelia interaction. **(F)** Quantification of cortical CCL2 levels in the uninjured and stroked (1 dpi) brains from WT and Slc4a4-icKO mice. Data are presented as mean ± SEM. N = 3-5 mice per genotype. *p<0.05, **p<0.01 by Student’s t-test. **(G)** Quantification of astrocytic CCL2 expression from double immunostaining. Data are presented as mean ± SEM. n = 7-17 cells collected from N = 3-5 mice per genotype. *p<0.05 by Student’s t-test. **(H)** Quantification of endothelial CCR2 expression from double immunostaining. Data are presented as mean ± SEM. Each dot represents an individual animal. N = 4-5 per genotype. *p<0.05, *p<0.05 by Student’s t-test. **(I)** At 4 dpi, astrocytic CCL2 expression was visualized by double immunostaining of CCL2/GFAP. Endothelial and astrocytic CCR2 expression was visualized by double immunostaining of CCR2/CD31 and CCR2/Aldh1l1-GFP respectively.

To further demonstrate the putative role of the Slc4a4-CCL2-CCR2 axis in astrocyte-endothelial crosstalk, we perform *in vivo* rescue experiments using an established antibody known to block CCL2 function (FBα) (44, 45) and an established CCR2 antagonist RS504393 (46) (Figure 6A). We treated a mouse endothelial cell line with WT-CM and KO-CM in combination with CCL2 FBα and examined the paracellular pathway by staining for tight junctional markers ZO-1 and Claudin-5. CCL2 inhibition was found to restore junctional marker expression (Figure 6, B-D). We then performed a TEER assay to assess paracellular permeability and found that CCL2 inhibition restored KO-CM-induced paracellular leakage of cultured endothelial cell monolayers (Figure 6, E and F) to control levels. Furthermore, we investigated whether the astrocytic Slc4a4-CCL2 axis regulates endothelial transcellular pathways by performing cell uptake assays with fluorescence-conjugated albumin (Alb) and transferrin (Tf) as indicators for caveolae- and clathrin-mediated endocytosis, respectively. We found that KO-CM increased endothelial uptake of Alb but not Tf compared with WT-CM (Figure 6, G and H), suggesting that loss of astrocytic Slc4a4 promotes caveolae-mediated transcellular transport. Western blot analysis of CM-treated endothelia revealed increased pCav-1 expression upon KO-CM treatment, confirming the activation of caveolae-mediated transport, which was rescued by CCL2 inhibition (Figure 6, I and J). Consistent with our *in vivo* observations that loss of astrocytic Slc4a4 upregulates endothelial CCR2 (Figure 5, H and I), treatment of KO-CM increased endothelial CCR2 expression *in vitro*, which was rescued by CCL2 FBα (Figure 6I **and Supplemental Figure 7C**). Lastly, we blocked endothelial CCR2 function using the CCR2 antagonist RS504393 during the incubation with WT- or KO-CM and found that blockage of endothelial CCR2 restored both increased paracellular (ZO-1) and transcellular (pCav-1) pathways in KO-CM treated endothelia (Figure 6, K and L). Together, these findings indicate that loss of astrocytic Slc4a4 promotes paracellular and transcellular leakage by enhancing the astrocytic CCL2-endothelial CCR2 signaling axis (Figure 6M).

**Figure 6.**
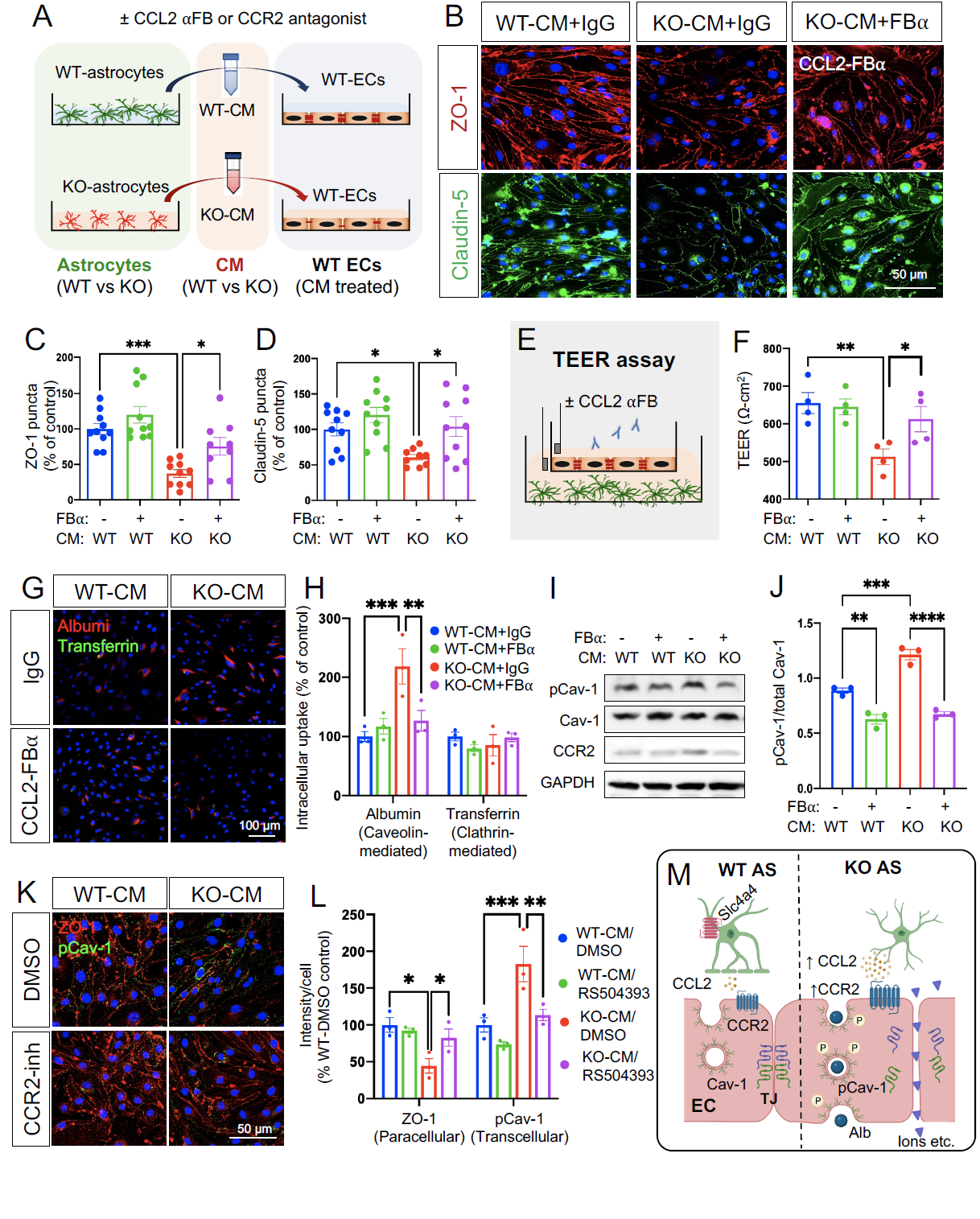
Slc4a4 regulates astrocyte-endothelia interaction via CCL2-CCR2 axis. **(A)** Mouse endothelial cells (bEnd3) were incubated with conditioned media (CM) collected from primary WT and Slc4a4 KO astrocytes with CCL2 functional blocking antibody (CCL2 ⍺FB, 15 ng/ml) or CCR2 antagonist for 24 hours. **(B-D)** Representative immunofluorescence images and quantification of tight junction proteins (ZO-1 and Claudin-5) of bEnd3 cells incubated with WT- or Slc4a4 KO-CM in the presence of control IgG or CCL2-FB⍺. Data are presented as mean ± SEM. Each dot represents a well collected from 3 independent experiments. *p<0.05, ***p<0.001 by two-way ANOVA. **(E)** Experimental setup of the transendothelial electrical resistance (TEER) assay. A monolayer of bEnd3 cells was co-cultured with WT or Slc4a4-deficient astrocytes, where astrocytes were plated at the bottom of the well and endothelial cells were plated in the insert, allowing endothelial cells to be exposed to astrocytic secreting factors without physical contact with astrocytes. CCL2 blocking antibody was added into the insert with a final concentration of 15 ng/ml. **(F)** The electro-resistance of the endothelial cell monolayer was measured as an indicator for the permeability of endothelial cells in the TEER assay. Data are presented as mean ± SEM. Each dot represents each independent culture. N = 4 independent assays. *p<0.05, **p<0.01 by two-way ANOVA. **(G-H)** Caveolin- and clathrin-mediated endothelial intracellular uptake was examined by Texas Red conjugated albumin and A488-transferrin, respectively, in bEnd3 cells incubated with WT- or Slc4a4 KO-CM with CCR2 FBα or control IgG. Data are presented as mean ± SEM. Each dot represents each independent culture. N= 3 independent assays. **p<0.01, ***p<0.001 by two-way ANOVA. **(I-J)** Western blot analysis of pCav-1, Cav-1 and CCR2 expression in bEnd3 cells incubated with WT- or Slc4a4 KO-CM with CCR2 FBα or control IgG. Data are presented as mean ± SEM. Each dot represents each independent culture. N = 3 independent assays. **p<0.01, ***p<0.001, ****p<0.0001 by two-way ANOVA. **(K-L)** Paracellular and transcellular endothelial transport in bEnd3 cells incubated with WT- or Slc4a4 KO-CM with CCR2 antagonist RS504393 (10 μM) or control DMSO were examined by immunostaining of ZO-1 and pCav-1. Data are presented as mean ± SEM. Each dot represents each independent culture. N = 3 independent assays. *p<0.05, **p<0.01, ***p<0.001 by two-way ANOVA. **(M)** A proposed mechanism by which astrocytic Slc4a4 regulates endothelial paracellular and transcellular transport pathways via the CCL2-CCR2 axis.

### Loss of CCL2 rescues BBB integrity in the absence of Slc4a4 after stroke

Our *in vitro* results implicated astrocytic CCL2 as a key downstream candidate of Slc4a4-mediated astrocyte-BBB interaction. We next examined the Slc4a4-CCL2 functional relationship *in vivo*. To this end, we intraperitoneally injected CCL2 FBα at 1 dpi, followed by BBB leakage assessment and marker analysis at 4 dpi (Figure 7A). Notably, CCL2 FBα treatment reversed BBB leakage (Evans blue, Fibrinogen) (Figure 7, B and C), increased CD31 expression (Figure 7, B and D), and rescued paracellular (Claudin-5) (Figure 7, B and E) and caveolae-mediated transcellular BBB leakage (pCav-1, Cav-1) (Figure 7, B and F; **Supplemental Figure 8**) in Slc4a4-icKO mice, without significant impact on reactive astrocytes at 4 dpi (GFAP) (Figure 7, B and G).

**Figure 7.**
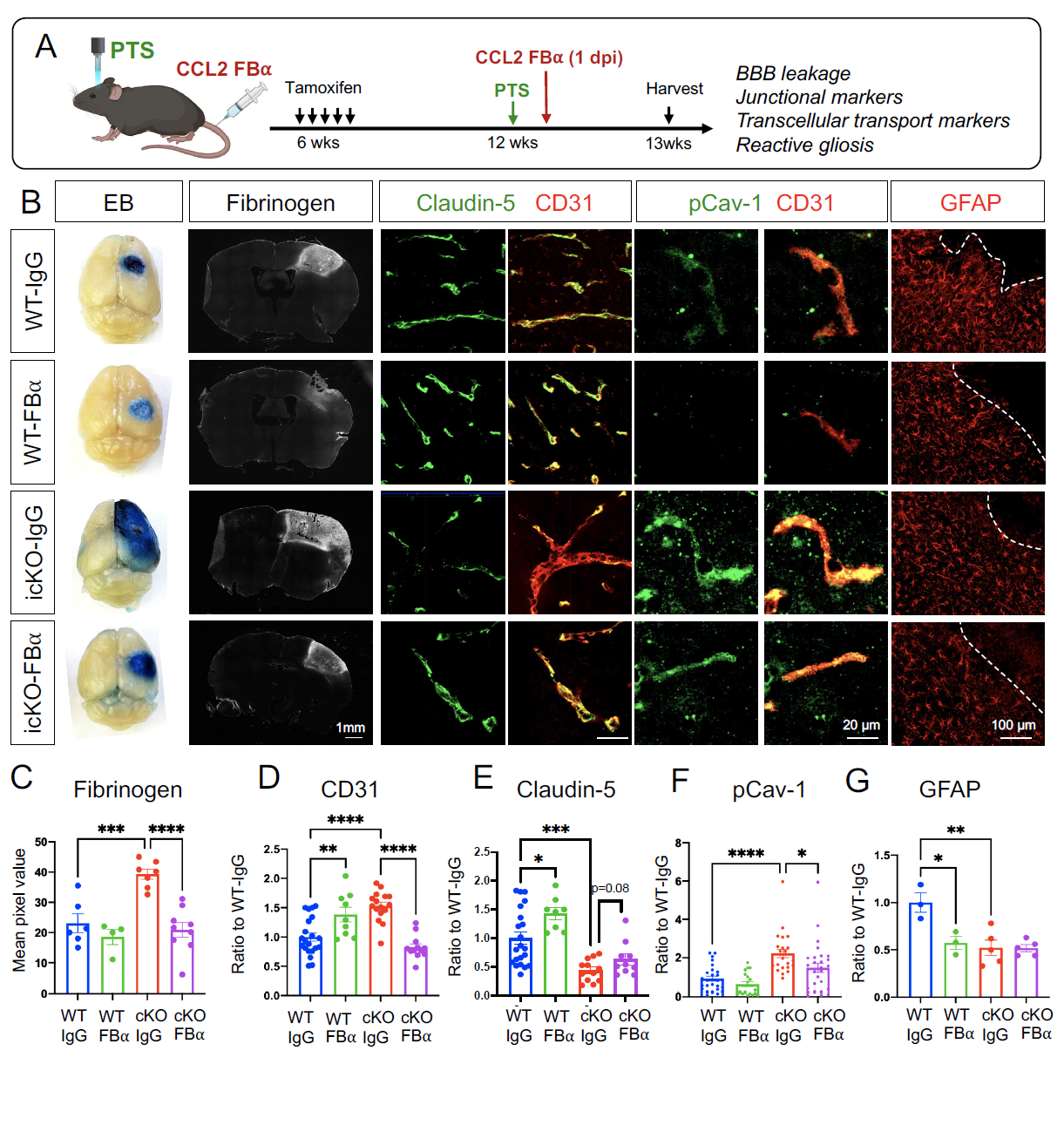
Pharmacologically CCL2 inhibition rescues loss of Slc4a4-induced exacerbated BBB damage after ischemic stroke. **(A)** Experimental scheme of the PTS induction in WT and Slc4a4-icKO mice, followed by intraperitoneal injection of CCL2 functional blocking antibody (CCL2-FB⍺) at 1 dpi. Brains were then harvested and analyzed at 4 dpi. **(B)** Representative images of protein leakage (Evans blue, fibrinogen), endothelial junctional marker expression (Claudin-5+; CD31+), endothelial pCav-1 expression and reactive astrocyte marker GFAP at the peri-lesion area in WT and Slc4a4-icKO mice with or without CCL2-FB⍺ treatment. **(C)** Quantification of fibrinogen intensity from immunostaining. Data are presented as mean ± SEM. Each dot represents an individual animal (N = 4-9 per group). ***p<0.001, ****p<0.0001 by two-way ANOVA. **(D)** Quantification of CD31 intensity from immunostaining. Data are presented as mean ± SEM. n = 2-3 blood vessels per animal and N = 4-5 animals per group. **p<0.01 ***p<0.001, ****p<0.0001 by two-way ANOVA. **(E)** Quantification of the intensity of Claudin-5 colocalized with CD31. Data are presented as mean ± SEM. n = 2-3 blood vessels per animal and N = 4-5 animals per group. *p<0.05, ***p<0.001 by two-way ANOVA. **(F)** Quantification of the intensity of pCav-1 colocalized with CD31. Data are presented as mean ± SEM. n = 4-5 blood vessels per animal and N = 4-5 animals per group. *p<0.05, ***p<0.001 by two-way ANOVA. **(G)** Quantification of GFAP intensity from immunostaining. Data are presented as mean ± SEM. Each dot represents an individual animal. N = 3-5 per genotype. *p<0.05 **p<0.001 by two-way ANOVA.

To further determine the putative role of the astrocyte-derived CCL2 in Slc4a4-mediated astrocyte-BBB interaction, we performed double loss-of-function studies in conditional knockout mice for both Slc4a4 and CCL2. As such, we generated temporally controlled conditional null alleles in the astrocytic lineage by crossing floxed alleles with the Aldh1l1-CreER line (Aldh1l1-CreER; Slc4a4^F/F^ denoted as Slc4a4-icKO, Aldh1l1-CreER; CCL2^F/F^ denoted as CCL2-icKO, and Aldh1l1-CreER; Slc4a4^F/F^; CCL2^F/F^ denoted as double-icKO) (Figure 8A). These mice were administered with tamoxifen at 5 weeks of age and Cre recombination/gene deletions were validated 5 weeks thereafter (**Supplemental Figure 9A**). We first examined BBB integrity by the expression of tight junctional markers in blood vessels. Notably, deletion of astrocytic CCL2 rescued Claudin-5 loss and blood vessel enlargement caused by the absence of Slc4a4 (**Supplemental Figure 9, B-D**), supporting an antagonistic relationship between Slc4a4 and CCL2 in the adult mouse brain under physiological conditions *in vivo*. To examine if the Slc4a4-CCL2 axis is conserved after ischemic stroke, we induced PTS 4 weeks after tamoxifen injection and analyzed the brain at 4 dpi. After stroke, deletion of astrocytic CCL2 rescued several phenotypes by Slc4a4 loss, including infarct size (Figure 8B **and Supplemental Figure 10A**), BBB leakage (Figure 8C **and Supplemental Figure 10A**), increased CD31 intensity (Figure 8, D and E; **Supplemental Figure 10, B and C**), impaired tight junctional marker expression (Claudin-5) (Figure 8, D and F; **Supplemental Figure 10, B and D**), and enhanced transcellular transport (pCav-1, Cav-1) (Figure 8, D and G; **Supplemental Figure 10, B and E**) surrounding the peri-lesion region. Importantly, genetic deletion of the astrocyte-specific CCL2 restored the number of vessel-associated astrocytes and vessel coverage by astrocytes in Slc4a4-icKO mice after stroke (Figure 8, D and H; **Supplemental Figure 10F**). These findings provide compelling evidence for the genetic hierarchy in which CCL2 functions downstream of Slc4a4 in BBB maintenance and reconstruction after stroke injury. Taken together, our results from both pharmacological and genetic approaches support the notion that Slc4a4 governs astrocyte-endothelial interaction by regulating astrocytic CCL2 under both physiological and stroke conditions.

**Figure 8.**
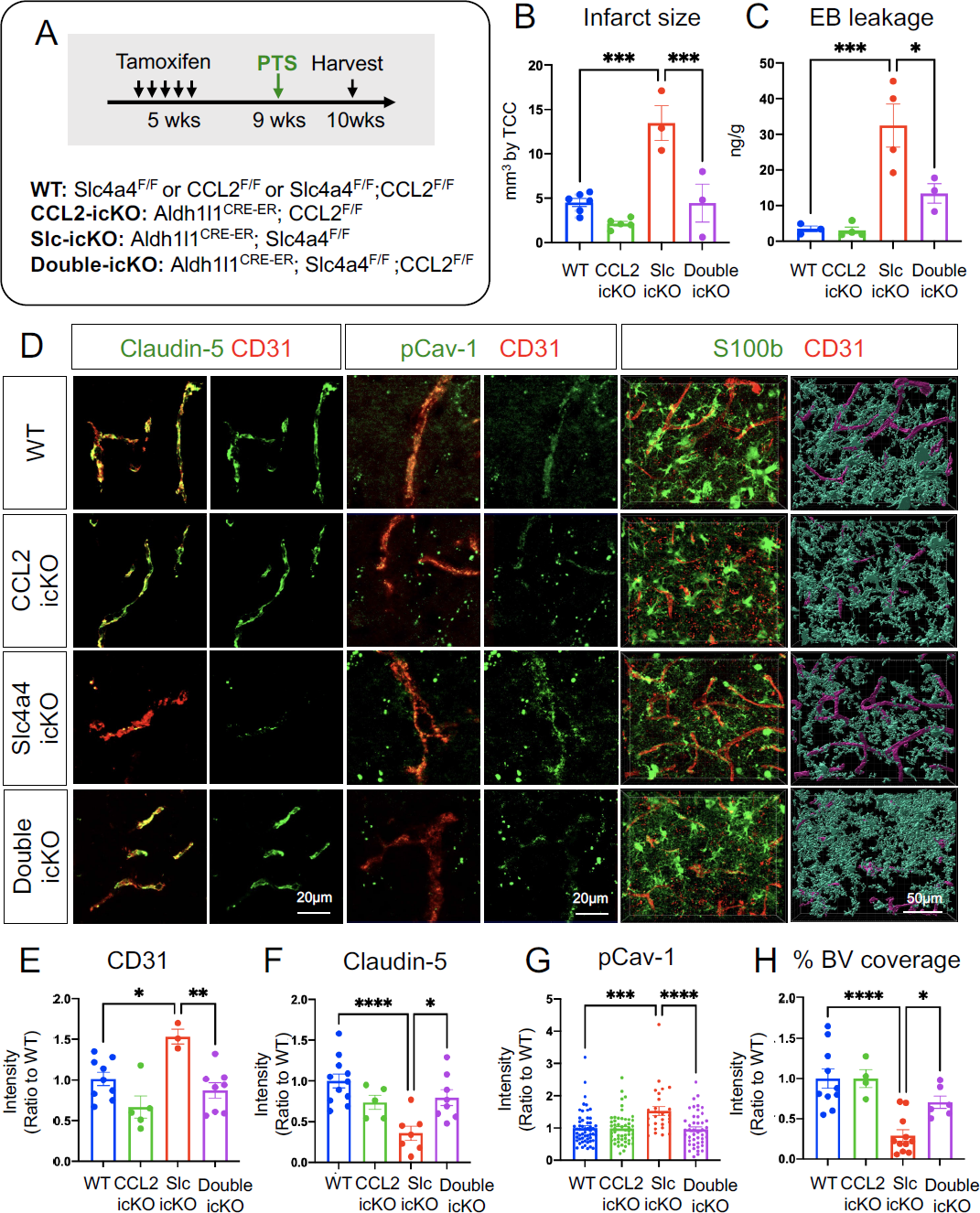
Genetic inhibition of astrocytic derived CCL2 rescues loss of Slc4a4-induced exacerbated BBB damage after ischemic stroke. **(A)** Experimental scheme of the PTS induction in temporally controlled astrocyte-specific conditional null alleles. Brains were then harvested and analyzed at 4 dpi. **(B)** Quantification of infarct size based on 2,3,5-Triphenyltetrazolium chloride staining of serial 1mm-thick brain sections from stroked brains at 4 dpi. Data are presented as mean ± SEM. Each dot represents an individual animal. N = 3-6 animals per group. ***p<0.001 by two-way ANOVA. **(C)** Evans blue levels in stroked brains from WT and Slc4a4-icKO mice were determined by colorimetric assays. Data are presented as mean ± SEM. Each dot represents an individual animal. N = 3-4 per genotype. *p<0.05, ***p<0.001 by two-way ANOVA. **(D)** Representative images of endothelial junctional marker expression (Claudin-5+; CD31+) and endothelial pCav-1 at the peri-lesion area. Reactive astrocyte and blood vessel interactions are visualized by double fluorescence staining of S100b and CD31. **(E)** Quantification of CD31 intensity at the peri-lesion area. Data are presented as mean ± SEM. n = 1-2 blood vessels per animal and N = 3-5 animals per group. *p<0.05, **p<0.01 by two-way ANOVA. **(F)** Quantification of Claudin-5 intensity colocalized with CD31 at the peri-lesion area. Data are presented as mean ± SEM. n = 1-2 blood vessels per animal and N = 3-5 animals per group. *p<0.05, ****p<0.0001 by two-way ANOVA. **(G)** Quantification pCav-1 intensity colocalized with CD31 at peri-lesion area. Data are presented as mean ± SEM. n = 5-6 blood vessels per animal and N = 5-6 animals per group. ***p<0.001, ****p<0.0001 by two-way ANOVA. **(J)** Quantification vessel area covered by astrocytes using IMARIS 3D reconstruction at the peri-lesion area. Data are presented as mean ± SEM. n = 1-2 images per animal and N = 4-5 animals per group. *p<0.05, ****p<0.0001 by two-way ANOVA.

### Slc4a4 regulation of CCL2 secretion is mediated by arginine metabolism

We next sought to decipher the mechanism by which Slc4a4 regulates astrocytic CCL2 secretion. Given the established link between astrocyte-mediated pH regulation and cellular metabolism, we performed unbiased metabolomic profiling in the cortex of the Slc4a4-icKO and WT mice. Arginine metabolism was identified as one of the top altered pathways (**Supplemental Figure 11A**). Specifically, we observed increased conversion of arginine to both citrulline and polyamine metabolites, resulting in reduced levels of arginine and increased levels of citrulline, putrescine and spermidine in Slc4a4-icKO mice compared with WT controls (Figure 9, A and B). Interestingly, Slc4a4-icKO mice also exhibited an increase in nitric oxide (NO), a byproduct of arginine to citrulline metabolism (Figure 9C), and a factor known to be critical for BBB breakdown as well as promoting proinflammatory cytokines production (47, 48).

**Figure 9.**
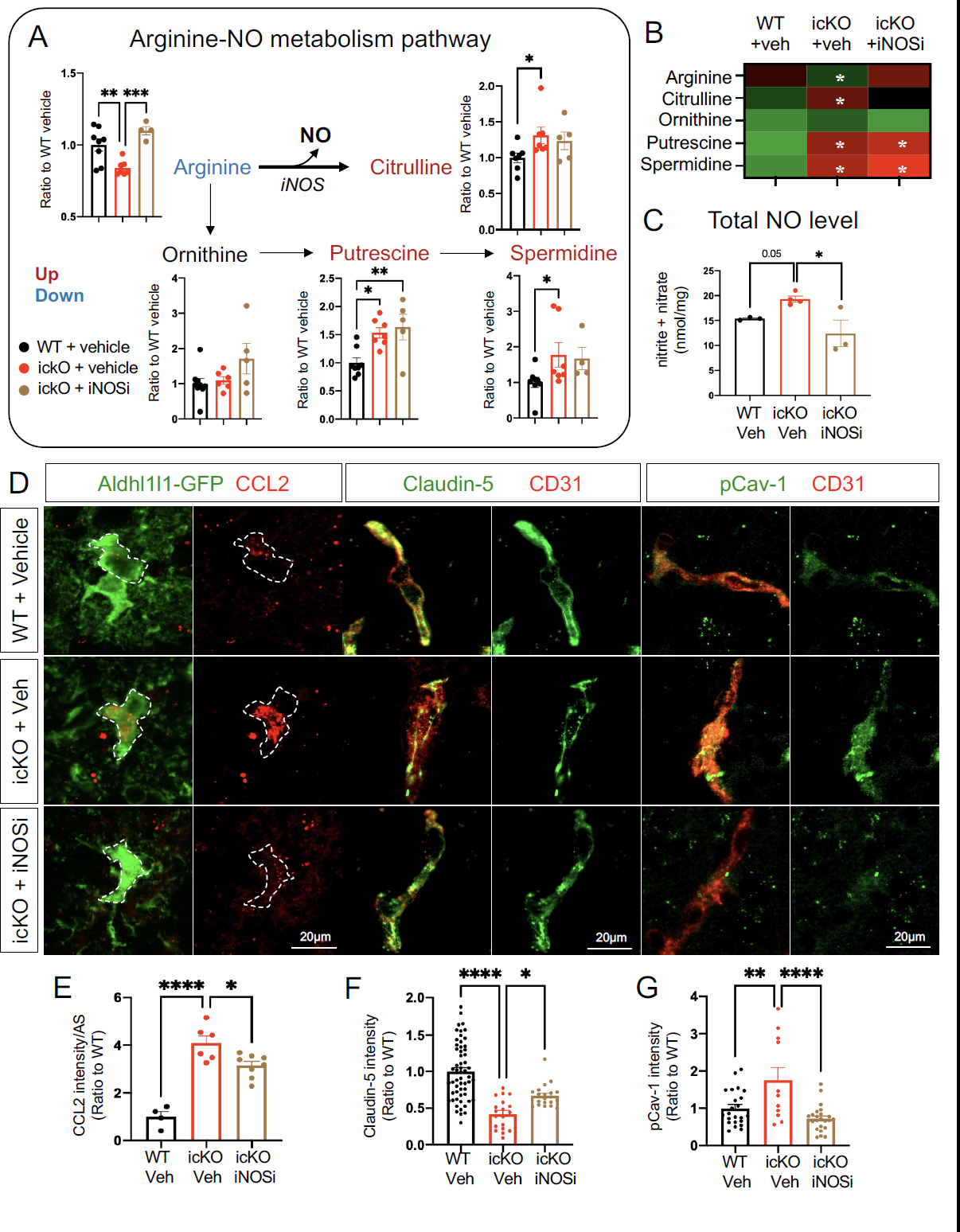
Astrocytic CCL2 dysregulation by the loss of Slc4a4 is partially attributable to arginine hypermetabolism. (**A-B**) WT and Slc4a4-icKO mice were intraperitoneally injected with iNOS inhibitor (L-NMMA, 10mg/kg) from 1-3 dpi. Brains were harvested at 4 dpi for analysis of cortical arginine metabolites. Data are presented as mean ± SEM. N = 4-7 animals per group. *p<0.05, **p<0.01, ***p<0.001 by one-way ANOVA. **(C)** Total NO levels in stroked cortices were measured by the total concentration of nitrite and nitrate using a colorimetric assay. *p<0.05 by one-way ANOVA. **(D-G)** Representative images and quantification of CCL2 colocalized with Aldh1l1-GFP, Claudin-5 colocalized with CD31, and pCav-1 colocalized with CD31 at the peri-lesion area. Data are presented as mean ± SEM. Each data point represents individual images from multiple animals. N = 3-4 animals per group. *p<0.05, **p<0.01, ****p<0.0001 by one-way ANOVA.

Given the known role of Slc4a4 in transporting arginine-rich peptides (49), we chose arginine metabolism as one of Slc4a4’s downstream pathways to further investigate if increased citrulline/NO production from arginine metabolism contributes to elevated CCL2 levels in the absence of astrocytic Slc4a4. To this end, we treated Slc4a4-icKO and WT mice with an inhibitor of NO synthesis, which targets iNOS, the rate-limiting enzyme of the arginine to citrulline metabolic reaction (**Supplemental Figure 11B**). As expected, the iNOS inhibitor rescued excess arginine to citrulline/NO conversion in Slc4a4-icKO brains, but not arginine to polyamine conversion (Figure 9, A-C). Although iNOS inhibition did not rescue the infarct size, it ameliorated BBB leakage at 4 dpi in Slc4a4-icKO mice (**Supplemental Figure 11, C-E**). Moreover, we observed a partial but significant rescue of CCL2 overproduction in Slc4a4-icKO mice after stroke by iNOS inhibition (Figure 9, D and E), along with restored expression of the tight junction marker Claudin-5 and amelioration of caveolae-mediated transcellular leakage (Figure 9, D, F and G). Together, these results provide important functional evidence that increased arginine to citrulline/NO conversion contributes to astrocytic chemokine secretion, consequently modulating endothelial cell integrity and BBB maintenance after ischemic stroke.

## Discussion

The mechanism underlying BBB maintenance and recovery after injury remains poorly understood. In this study, we discovered that Slc4a4, an astrocyte-enriched Na^+^/HCO_3_^−^ cotransporter, is required for BBB maintenance and reconstruction after ischemic stroke injury. Using both pharmacological and lineage-specific loss-of-function approaches, we provide compelling evidence that astrocytic Slc4a4 ablation stimulates chemokine CCL2 production, which in turn exacerbates BBB damage after stroke. Mechanistically, the over-production of CCL2 upon Slc4a4 loss is partially mediated by exaggerated NO production from arginine metabolism. Overall, our study reveals a novel molecular cascade where Slc4a4 regulates astrocyte-endothelial cell interaction via coupling of intracellular metabolism and the secretome, highlighting a novel therapeutic strategy to stimulate BBB recovery after ischemic stroke.

pH homeostasis affects every facet of brain activity, including neuronal transduction (24), cellular metabolism (15, 50) and immune response (51). Furthermore, pH dysregulation has been linked with several devastating neurological disorders, including ischemic stroke and brain cancers (52, 53). In the normal brain, many fundamental brain activities, ranging from neurotransmitter release to cellular metabolism, result in a rapid rise of extracellular H^+^. Pathological pH decline is often observed after ischemic stroke, partially owing to hypoxia, which then triggers a sharp response by ion transporters across multiple cell types (15). Recent studies showed that Slc4a4 is activated and releases HCO_3_^-^ to neutralize rapid acidification from synaptic transmission (24). In the current study, we provide evidence for a novel role of Slc4a4 in the astrocyte’s support for BBB function and recovery after ischemic stroke. Interestingly, previous studies reported that astrocytic Na^+^/H^+^ exchanger isoform 1 (NHE1) performs opposite functions in the context of ischemic stroke: loss of astrocytic NHE1 reduced astrogliosis and ameliorated BBB damage (20). It should be noted that contrary to Slc4a4’s function to neutralize acidic pH, NHE1 is known as a major mediator for H^+^ extrusion in astrocytes regardless of the extracellular pH (54). Unlike most pH regulators in the brain, which are either acid extruders or intruders, Slc4a4 transports sodium/bicarbonate bidirectionally across the cell membrane depending on the electrogenic gradients (55, 56), making it a versatile ‘pH modulator’ for drug-targeting in brain disorders.

Moreover, a previous study reported a subpopulation of cortical astrocytes expressing high levels of Slc4a4 and carbonic anhydrase 2 (24), suggesting potential crosstalk of Slc4a4 and intracellular carbonic metabolism. Yet, it is unknown if loss of astrocytic Slc4a4 would lead to compensatory changes in the expression or activity of pH regulators in astrocytes or other cell types (e.g., neurons, oligodendrocytes) under physiological conditions or after ischemic stroke. While our data clearly indicate that astrocytic Slc4a4 is critical for BBB maintenance and reconstruction, it would be important to delineate further whether astrocytic Slc4a4 function is also dependent on the expression/function of other pH regulators and/or enzymes.

Injury-induced BBB leakage has been linked with increased paracellular transport regulated by tight junction proteins and elevated caveolae-mediated transcellular transport (57). Mounting evidence suggests that the CCL2/CCR2 signaling is detrimental to ischemic stroke recovery and BBB function (58, 59). Increased CCL2 leads to loss of endothelial tight junction marker expression (60) and upregulation of caveolae (61). However, it is unclear how CCL2 secretion is regulated in a cell type-specific manner. Our study revealed Slc4a4 as a key regulator for astrocyte sourced CCL2. This discovery, coupled with results from our unbiased metabolic profiling and pharmacological inhibition, supports the model that Slc4a4 regulates astrocytic CCL2 secretion partially via global arginine/NO metabolism. Several types of cells in the brain can produce NO. On one hand, endothelial cells utilize endothelial nitric oxide synthase (eNOS) to produce NO at physiological states, which is essential for maintaining normal endothelial function (62). On the other hand, several other types of cells, including astrocytes and microglia, utilize iNOS to produce extra NO under injury, which is known to play a detrimental role in BBB function (63). While our study shows that loss of astrocytic Slc4a4 stimulates iNOS-dependent NO production, further studies will examine the expression and activity of key enzymes involved in brain arginine metabolism to define the arginine/NO-CCL2 relationship in a cell type-specific manner.

Our results also indicate that Slc4a4 plays an intrinsic role in astrocytogenesis during development as well as reactive gliosis after ischemic stroke injury. These findings underscore the mechanistic convergence between glia development and regeneration after injury. While such conserved functions have been shown for key astrocytic transcriptional factors (64), our study reveals that intracellular or extracellular microenvironment factors are also critical for astrocytogenesis at a non-transcriptional level. Due to the technical limitation of direct measurement of intracellular pH at single-cell resolution, one remaining question from our study is by which mechanism Slc4a4 regulates astrocyte generation during development and after injury. Since metabolic diversity is one of the major factors driving reactive astrocyte response and heterogeneity in neurological disorders (65), it is plausible to speculate that Slc4a4 regulates astrocytogenesis by influencing pH-dependent cellular metabolism required for astrocyte development and reactive astrogliosis after stroke. Despite growing evidence showing that astrocytes are the powerhouse and recycling center in response to brain energy needs (6), less is known regarding the energy need and metabolic regulation intrinsic for astrocytes. It will be important to further decipher how Slc4a4 regulates gliogenesis, perhaps via a mechanism linked to intrinsic astrocytic cellular metabolism.

From the clinical perspective, patients carrying Slc4a4 variants show an increased risk for not only stroke (26–28), but also other CNS malfunctions, including migraine, epilepsy, and intellectual disability (29, 30). While pH/ion disturbance, BBB leakage and metabolic adaptation have been involved in the disorders mentioned above, whether Slc4a4-mediated astrocyte-BBB interaction plays a role in the other brain disorders remains to be investigated. Our study provides an important mechanistic entry point to dissect the role of Slc4a4 in other CNS malignancies.

## Material and methods

### Animals

All mice were maintained and studied according to protocols approved by the Institutional Animal Care and Use Committee of Baylor College of Medicine. All genotypes were on the C57BL/6J strain and blinded for data acquisition and analysis and experiments included both genders and age-matched controls. Aldh1l1-EGFP transgenic mice were used for astrocyte-specific labeling (66). For temporal expression of Cre in astrocytes, Aldh1l1-Cre^ERT2^ (The Jackson Laboratory, stock number 029655) was used. Slc4a4-floxed mice were generated and provided by Dr. Shull (67). CCL2-floxed animals were purchased from The Jackson Laboratory (stock number 029655). To induce gene deletion during development, animals were injected subcutaneously with tamoxifen (Sigma T5648, 100 mg/kg, dissolved in corn oil) at postnatal day 3 (P3). To induce gene deletion in adult mice, animals were injected intraperitoneally with tamoxifen (100 mg/kg) daily for 5 consecutive days from 5 to 6 weeks old.

### FACS sorted astrocytes for RT-qPCR

Cortex was microdissected and dissociated into a single-cell suspension in FACS buffer (Leibowitz Media, supplemented with 1 mg/ml BSA, 10 mM HEPES pH 7.05, pen/strep, and 25 μg/ml DNaseI). For cell sorting, a BD FACSAria III instrument (100-μm nozzle and 20-p.s.i. setting) was used with FACSDIVA software. Around 100,000 GFP+ astrocytes were collected directly into 500 μl of Buffer RLT (QIAGEN 79216) with 1% β-Mercaptoethanol. Finally, each sample was vortexed and rapidly frozen on dry ice. RNA was extracted from pelleted cells using RNeasy Micro Kit (QIAGEN 74004), followed by cDNA synthesis (AmfiSure ONE PCR Master Mix P7000-050) and RT-qPCR (AmfiRivert cDNA Synthesis Platinum Master Mix, R5600) on a BioRad CFX Duet Real-Time PCR System. qPCR was carried out at 95°C for 30 s, 40 cycles of 95°C for 5 s and 60°C for 30 s, with subsequent melting curve analysis. Expression of transcripts of target genes was normalized to Gapdh. Primer sequence for mouse Slc4a4 used was obtained from published literature (68).

### Photothrombotic focal ischemia model

Focal ischemic sites were generated in the cortex of 10 to 12-week-old mice. Briefly, mice were anesthetized using isoflurane, and an incision was made to expose the skull. A customized dark PVC mask with a 2 mm diameter aperture was placed over the exposed skull to limit the laser illumination to a location 1 mm anterior and 1 mm lateral to bregma. Then mice were injected intravenously with 20 mg/kg of Rose Bengal dye (Sigma 330000, 10 mg/ml in sterile saline), followed by immediate exposure to light from Leica KL 300 LED for 15 minutes. The surgical site was then sutured, and the animals were allowed to recover in sternal recumbency until fully awake. For the pharmacological CCL2 rescue experiment, mice were intraperitoneally injected with CCL2 functional blocking antibody (R&D; AF-479-NA, 0.5 mg/kg) at 1 dpi; normal goat IgG (R&D; AB-108-C) was used as control. For the inhibition of iNOS, mice were intraperitoneally injected with L-NMMA (Sigma M7033, 10 mg/kg) once daily from 1 to 3 dpi; saline was used as a control. Upon harvesting of the tissue at designated time points, the animals were perfused, and tissues were processed as described below. For histology analysis, the lesion border is identified based on the autofluorescence signal and brightfield tissue histology and the peri-lesion is defined as 150μm outside the border. GFAP immunostaining was used to identify glial scar, which is defined by a dense limiting border consisting of fibrous reactive astrocytes surrounding the core lesion of stroke as shown in Supplemental Figure 5A. The thickness of the glial scar is measured as the maximum distance between the dorsal and the ventral side of the glial scar. For CCL2 quantification, metabolomic profiling and NO measurement, fresh brain cortical tissue was used and stroked tissue was microdissected based on white opaque scar-like appearance.

### TTC staining

Within 3 min of sacrifice, stroked brains were removed and sectioned into 1mm-thick brain slides using a brain matrix (Ted Pella). Brain slices were then immersed in a 2% solution of TTC in normal saline at 37°C for 30 min, after which sections were fixed in 4% PFA for photography.

### BrdU labeling *in vitro* and *vivo*

To evaluate endothelial cell proliferation in vitro after conditioned medium treatment, 10μm of BrdU was added to cultured cells 6hr before collecting. To evaluate astrocyte proliferation *in vivo* after stroke, we administered multiple doses of BrdU (200mg/kg, every 12 hours from 1-3 dpi) before harvesting. Brains were harvested and processed as described above. For the BrdU staining, an additional 2N HCl treatment for 30 minutes was performed.

### *In situ* hybridization, immunofluorescence staining and H&E histology

For immunofluorescence staining of tight junction makers (ZO-1, Claudin-5) and CCL2, brains were emersed in OCT and then snap-frozen on dry ice and stored at −80 °C until ready for sectioning. Tissue sections were then prepared with a cryostat (Leica) at a thickness of 15μm, followed by 95% ethanol fixation. For *in situ* hybridization and immunofluorescence staining, brains were fixed in 4% paraformaldehyde (PFA) overnight, dehydrated in gradient sucrose, embedded in OCT blocks (Tissue TEK), and stored at −80°C until ready for sectioning at a thickness of 15 to 100 μm.

For *in situ* hybridization, RNA probes of mouse Slc4a4 with DIG and/or FITC labels were generated in-house to detect the corresponding RNAs. All probes were tested for specificity and sense probes were included in the experiments as a control. Detailed procedures and reagents were described before (69).

For immunofluorescence staining, tissue slides were washed in PBS, permeabilized with PBST (0.3% Triton in PBS), blocked with 10% normal goat serum in PBST, and then incubated with primary antibodies overnight at 4°C, followed by secondary antibody and nuclear staining with DAPI. The following commercial primary antibodies were used: rabbit anti-GFAP (1:1,000; Agilent Dako), mouse anti-NeuN (1:500; Millipore), rabbit anti-GFP (1:1,000; Chromotek; PABG1-100), rabbit anti-NF1A (gift from Dr. Benjamin Deneen), rabbit anti-Sox9 (1:1,000; Millipore; AB5535), rat anti-BrdU (1:1,000; Abcam; ab6326), rabbit anti-S100b (1:1,000; Agilent Dako; Z0311), Rat anti-CD31(1:200; BD Bioscience; 55047), rabbit anti-Glut1 (1:500; Millipore; 07-1401), rabbit anti-AQP4(1:500; Sigma; A5971), rabbit anti-ZO-1 (1:250; Invitrogen;40-2200), Alexa 488 conjugated Claudin-5 (1:500; Invitrogen, 35-358-8), rabbit anti-CCL2 (1:500, PerproTech, 500-P113), goat anti-CCL2 (1:500; R&D, AF-479), ), rabbit anit-CCR2 (1:500; Abcam; ab273050), mouse anti-Cav-1 (1:500; BD Bioscience; 610406), mouse anti-pCav-1 (1:500; BD Bioscience; 611339). Secondary antibodies conjugated with Alexa fluor 488 and 568 (Thermo Fisher Scientific) were used at 1:1,000 dilution. Stained slides were mounted with VectaShield Antifade mounting medium. Images were obtained by Zeiss Imager.M2m equipped with ApoTome.2 or Leica STED microscope system.

For H&E staining, hematoxylin was stained for 5 minutes, then rinsed with tap water until no more color changes. 2–3 dips in the acidic solution (1% HCl in 70% EtOH) reduced the background. After rinsing with tap water, Eosin was stained for 1 minute, followed by tap water rinsing. Finally, dehydration was accomplished by three 5-minute incubations with 95% EtOH, 100% EtOH, and Xylene. Stained slides were mounted with Permount mounting medium. Images were obtained using a Nikon spinning disc microscope.

### 3D image and lightsheet confocal

To label blood vessels *in vivo*, tomato-labeled lectin (Lycopersicon Esculentum) (VECTOR Laboratories, #DL-1178-1) was injected into mice via tail vein (50 μl) and intracardially (100 μl) before harvesting. Astrocytes were genetically labeled with Aldh1l1-GFP. Mice were then perfused, and brains were fixed as described above. As described previously (70), brains were then cleared using the X-CLARITY platform (Logos Biosciences), followed by imaging using Lightsheet Z.1 microscope (Zeiss).

### Astrocyte morphology and astrocyte-blood vessel coverage analysis

For all morphological analysis by 3D rendering, astrocytes (Aldhl1l-GFP) or blood vessels (lectin) were randomly picked and analyzed by a person blinded to the experimental groups (H.O.) Morphological analysis was performed using the Filament module in the IMARIS software 10.3.0 (Oxford Instruments). Maximum orthogonal projections of Z-stack images (1 μm interval, 40-80 slides) were imported to IMARIS and astrocyte branches were outlined using the Filament Autopath algorithm with a dendrite starting point diameter of 5.00 μm and a dendrite seed point diameter of 0.216 μm. Following automated tracing, Filament structures were then manually pruned to eliminate false positive branches. For astrocyte-blood vessel 3D rendering, the fluorescence signal was reconstructed with the surface tool, and the area of a blood vessel (lectin) covered by astrocytes (Aldh1l1-GFP) was calculated by the software.

For astrocyte volume analysis, late P0/early P1 pups were sedated by hypothermia until anesthetized. 1 μL of AAV-PhP.eB -gfaABC1D-mCherry-CAAX construct (titer∼1 × 10^12^ GC/ml) mixed with Fast Green Dye was injected into the lateral ventricle of one hemisphere using a pulled glass pipette. Pups were recovered on a heating pad, returned to their home cage, and monitored until tissue collection at designated ages. To assess the volume of individual astrocytes in the L 4-5 visual cortex, 100 μm-thick floating sections containing astrocytes labeled sparsely with mCherry-CAAX were used. High magnification images containing an entire astrocyte (60-80 μm Z-stack) were acquired by Zeiss Imager.M2m equipped with ApoTome.2 with the 63x objective. Astrocytes in which the entire astrocyte could not be captured within the section were excluded. Astrocyte volume was calculated using Imaris Bitplane software with the surface tool.

### Calcium imaging

Calcium imaging of cortical astrocytes was performed as previously described (71). Briefly, AAV2/9-GfaABC_1_D-GCaMP6f virus (Addgene #52925) was stereotaxically injected into the cortex 2 weeks before imaging. On the day of imaging, animals were anesthetized with isoflurane, followed by intracardial injection of tomato-labeled lectin (Lycopersicon Esculentum) (VECTOR Laboratories, #DL-1178-1) to label blood vessels. The brain was isolated and sliced quickly. Calcium activity was collected at 1 Hz for 200s using a two-photon resonant microscope (LSM 7MP, Zeiss) with a Coherent Chameleon Ultra (II) Ti-sapphire laser tuned to 900 nm. Relative changes in fluorescence of soma, primary branches, and endfeet Relative changes in fluorescence of soma, primary branches, and endfeet were quantified using GECIquant with ImageJ. MATLAB software was used to quantify amplitude and frequency.

### BBB leakage assessment *in vivo*

Albumin leakage into the brain was assessed by Evans blue. Briefly, mice were injected intraperitoneally with a filtered 2% Evans blue solution in PBS (15 ml/kg). Mice were then perfused with PBS, and brains were harvested 24 hours (non-injured animal) or 6 hours (stroke animal) after injection. Brains were weighed and incubated with formamide (2 v/w) for 24 hours at 65°C. Subsequently, the samples were briefly centrifuged, and supernatants were taken to measure absorbance at 600 nm with a standard curve of Evans blue dye. The final concentration of Evans blue in the tissue was calculated as ng/g tissue.

3 kDa dextran leakage into the brain was assessed as previously described. Mice were intravenously injected with 3 kDa A488-dextran (10 mg/kg, ThermoFisher Scientific D3306). After 2 hours, mice were anesthetized and perfused with ice-cold PBS (pH 7.4) and PFA. Brains were harvested, PFA-fixed, dehydrated, sectioned and imaged as described above. The leakage of injected dextran into brain parenchyma was measured based on intensity and quantified using ImageJ software.

Small molecule leakage into the brain was assessed by EZ-Link Sulfo-NHS-Biotin (MW = 443 kDa, Thermo Fisher Scientific). Briefly, mice were intracardially injected with 200 μl biotin (20 mg/ml) right before harvesting. Brains were snap-frozen, followed by sectioning and ethanol fixation, as described above. Sections were stained with Alexa 535-streptoavidin (Invitrogen, 1:1,000), mounted and imaged with a Nikon spinning disc microscope. The leakage of injected biotin into brain parenchyma was quantified using ImageJ software.

### Western blot

Cells were lysed with lysis buffer (150 mM NaCl, 50 mM Tris–HCl, 1 mM EDTA, 1% Triton X-100, 1% NP-40) containing 1X protease inhibitor cocktail (GenDEPOT, P3100) and electrophoresed on 10% SDS-PAGE (10 µg protein) and transferred to nitrocellulose membranes. After blocking with 5% nonfat milk, the membranes were incubated with primary antibodies, including mouse anti-Cav-1 (1:500; BD Bioscience; 610406), mouse anti-pCav-1 (1:500; BD Bioscience; 611339), rabbit anit-CCR2 (1:500; Abcam; ab273050), and rabbit anti-Gapdh (1:5000, GeneTex, GTX100118) as a loading control. Blots were incubated with HRP-conjugated secondary antibodies. Immunoreactivity was developed using ECL reagent (GenDEPOT, W3653) and imaged with Bio-Rad Gel Imager ChemiDoc E.

### Extracellular pH after ischemic stroke

Mice with ischemic stroke were intraperitoneally injected with 100 μl pHLIP-ICG probe (1 mg/kg, synthesized and provided by Y.K.R.). 24 hours after injection, brains were harvested and imaged using the Bruker Xtreme Imager with 735 nm excitation and 830 nm emission wavelength. The fluorescence intensities in regions of interest were calculated using Image J.

### Cytokine array

Astrocyte-conditioned media was collected from 3 different mice, pooled, and centrifuged for 10 minutes at 1,000 rpm, and immediately divided into aliquots and stored at −80 °C until further analysis. Protein concentrations were determined by Bradford assay and adjusted to the same concentration. Conditioned media was then measured in duplicates with Mouse Cytokine Array Panel A (ARY006, R&D Systems) according to the manufacturer’s instructions. Signal detection was performed using a ChemiDoc Imaging System (Bio-Rad, Hercules, CA, USA), and the signal was quantified using Image Lab software (Bio-Rad).

### Metabolomics analysis

Mouse cortices were flash-frozen in liquid nitrogen and stored at −80 °C until analysis. Samples were subjected to unbiased metabolomic analysis as previously described (72).

Briefly, samples were homogenized by glass beads in a Bullet Blender (at setting 3 for 5 minutes at 4 °C). Suspensions were then centrifuged and supernatants were collected for metabolomics analysis by UHPLC-MS analysis (Vanquish – Qexactive, Thermo Fisher), as described (73). Metabolite assignments and isotopologue distributions were performed using MAVEN (Princeton). Metabolic pathway analysis was performed using the MetaboAnalyst 5.0 package (www.metaboanalyst.com).

### NO measurement

NO concentration was measured by Griess methods (Nitrate/Nitrite Colorimetric Assay Kit; Cayman Chemical Co; 780001) according to the manufacturer’s instructions.

### Primary astrocyte culture and *in vitro* BBB leakage assay

Astrocytes were obtained from the cortices of P0 to P1 WT and Slc4a4-cKO (Slc4a4^F/F^; Aldh1l1-cre) mice and cultured in DMEM/F12 medium containing 10% FBS and 1% penicillin-streptomycin. Cells were then plated onto poly-D-lysine coated T75 culture flasks, and media were changed every 3-4 days. After 7 days, flasks were placed on an orbital shaker at 120 rpm overnight to remove microglia and oligodendrocyte precursors.

For conditional media collection for cytokine array and mass-spec profiling, astrocytes were subcultured into 12-well plates and allowed to grow until confluent. 24 hours before collection, the medium was switched to a medium free of phenol and FBS. For the oxygen-glucose deprivation (OGD) model, the prepared cells were cultured in glucose-free Dulbecco modified Eagle medium (GenDEPOT) in a 37°C incubator containing a humidified gas mixture of 1% O_2_ / 95% N_2_ / 4%CO_2_ for 2 hours. For reoxygenation, the cells were incubated for 24 hours with normal DMEM containing 1% FBS in a 37°C incubator containing humidified 5% CO_2_ / 95% air.

For in vitro TEER assay, astrocytes were seeded at the bottom of the transwell plate (0.4-μm pore size, 12-well; Corning) at a density of 3 × 10^5^ cells per filter. After allowing the astrocytes to grow for 48 hours, immortalized mouse endothelial bEnd3 cells (American Type Culture Collection) were plated in the inserts (5 ×10^4^ cells per well). TEER measurements were then made 24 hours after bEnd3 seeding using an EVOM resistance meter (World Precision Instruments) with Stx2 electrodes. TEER resistance was measured in ohms after subtracting resistance from blank transwell inserts.

For in vitro endocytosis assay, bEnd3 cells were incubated with 50 μg/mL Alexa 488-transferrin (ThermoFisher Scientific, T13342) or Texas Red-albumin (ThermoFisher Scientific, A13100) for 30 min at 37°C, 5% CO_2_, as described previously (74). Cells were then rinsed three times with ice-cold PBS, fixed with 4% paraformaldehyde (PFA) and stained with DAPI. The samples were imaged and analyzed with Zeiss Imager.M2m equipped with ApoTome.2.

### Mass-spec protein profiling

Conditioned medium was subjected to unbiased mass-spec profiling as previously described (71). Briefly, samples were denatured and digested, followed by extraction with 50% acetonitrile and 2% formic acid. Vacuum-dried peptides were subjected to reverse phase column chromatography with a micro-pipette tip C18 column and fractionated with stepwise ACN gradient into different elution groups. Eluent was pooled and vacuum-dried for analysis with a nano-LC 1000 coupled with an Orbitrap Fusion^TM^ mass spectrometer (ThermoFisher Scientific). Data were analyzed with Proteome Discoverer 2.1 interface (ThermoFisher Scientific) and detected peptides were assigned into gene products by Py Grouper and Tackle analysis flatform from iSpec(75).

### Statistics

GraphPad Prism v8 was used for generating graphs and statistical analysis. Data were reported as mean ± SEM. Significance was calculated using two-tailed, unpaired Student’s t-tests for two-group comparisons, or two-way ANOVAs for four-group comparisons, followed by Sidak’s post hoc analysis. *P < 0.05, **P < 0.01, P*** < 0.001, and ****P < 0.0001 were considered to indicate statistical significance. Sample size for each experiment is indicated in the corresponding figure legend. Number of biological replicates, either animals (*N*) and/or cells/images (*n*), used for quantifications of each experiment were indicated in the respective figure legends.

## Acknowledgments

We would like to thank Diego Cortes for technical support in animal experiments. This work was supported by grants from the NIH/NINDS (R01NS110859 and R01NS126287 to H.K.L), National Multiple Sclerosis Society (RG-1907-34551 to H.K.L), Helis Medical Research Foundation, the Mark A. Wallace Endowment established by an anonymous donor (to H.K.L) and NIH/GMS (R01GM073857 to Y.K.R). This work was also supported by the Microscopy Core facilities (P50HD103555), Mouse Metabolism and Phenotyping Core (UM1HG006348, R01DK114356, R01HL130249), Optical Imaging & Vital Microscopy (OiVM) core and Mass Spectrometry Proteomics Core at Baylor College of Medicine.

## Author contribution

Q.Y and H.K.L conceived the project and design the experiments; Q.Y, J.J, C.W, H.O, T.J.C performed the experiments; Y.K.R, A.D.A, S.Y.J, C.Z, H.K.L provided reagent; S.P.M provided expertise. Q.Y, T.J.C and H.K.L wrote and edit the manuscript.

## Conflicts of interests

Y.K.R. is the founder of pHLIP Inc., but the company did not fund any part of the work reported here. The authors declare no other competing interests.

## Notes

### Competing Interest Statement

The authors have declared no competing interest.

