## supplementary figures for "Astrocytic Slc4a4 regulates blood-brain barrier integrity in healthy and stroke brains via a NO-CCL2-CCR2 pathway"

### **Supplemental Figure 1** Related to Figure 1

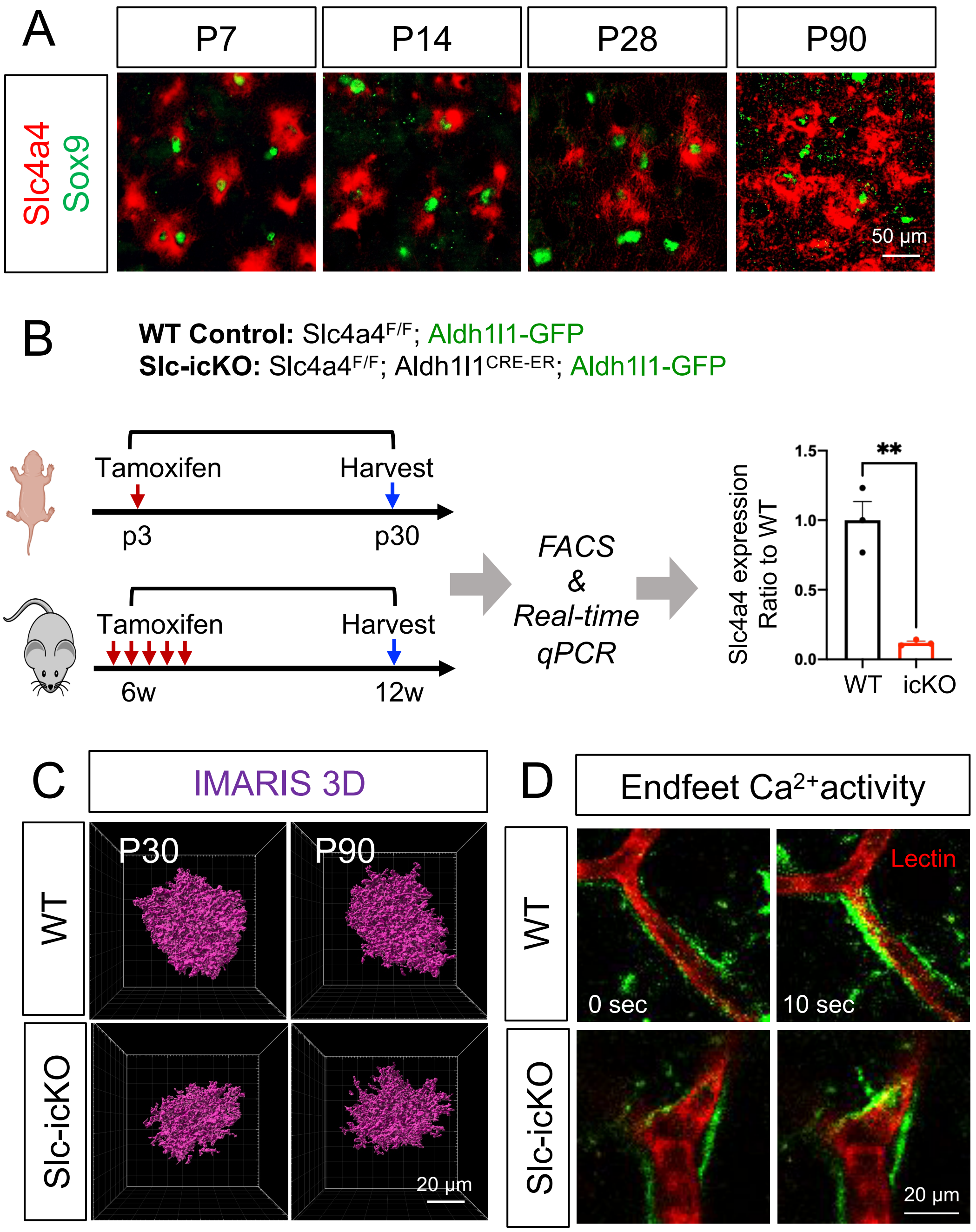


**Supplementary Figure 1. Slc4a4 expression during development and confirmation of Slc4a4 deletion in Slc4a4-icKO. (A)** Double in situ-immunofluorescence staining of Slc4a4 in astrocyte lineage (Sox9) in developmental and adult mouse cortex **(B)** Schematic of the generation of astrocyte-specific Slc4a4 knockout mice with Aldh1l1-GFP reporter. Deletion of Slc4a4 deletion was confirmed by quantitative RT-PCR of FAC-sorted GFP+ astrocytes. Data are presented as mean ± SEM. N = 3 per genotype. *p<0.05 by Student's t-test. **(C)** IMARIS 3D reconstruction of astrocytes sparsely labeled with AAV-GfaABC1D-mCherry-CAAX. **(D)** Representative images of astrocytic endfeet spontaneous calcium activity from WT and Slc4a4-icKO mice.

### **Supplemental Figure 2** Related to Figure 2

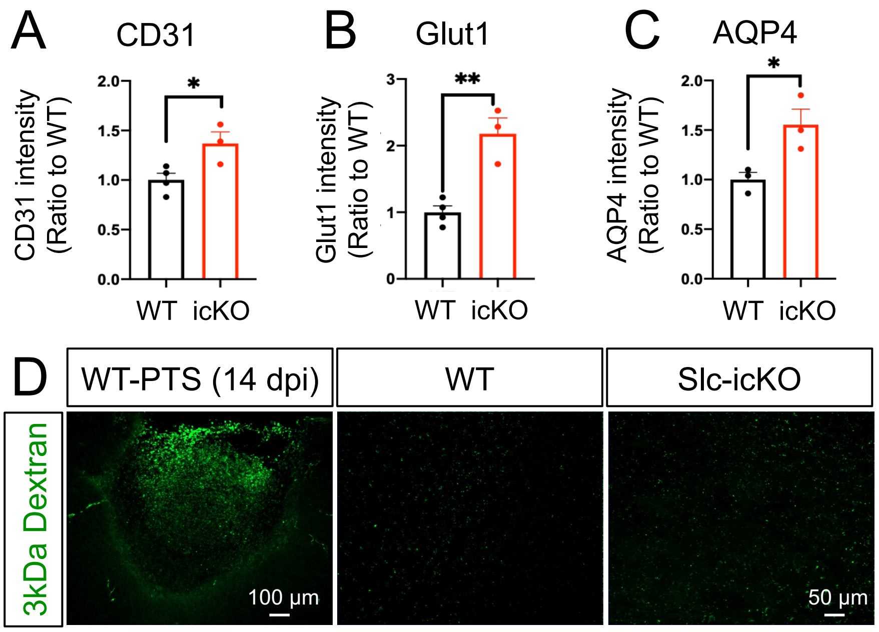


**Supplementary Figure 2. Loss of Slc4a4 increases the expression of endothelial markers but does not induce leakage of 3kDa dextran. (A-C)** Quantification of endothelial markers (CD31, Glut1, AQP4) based on intensity by immunostaining. **(D)** Representative images of brain sections after 3kDa FITC-dextran injection. Stroked brains were served as a positive control.

#### **Supplemental Figure 3** Related to Figure 3

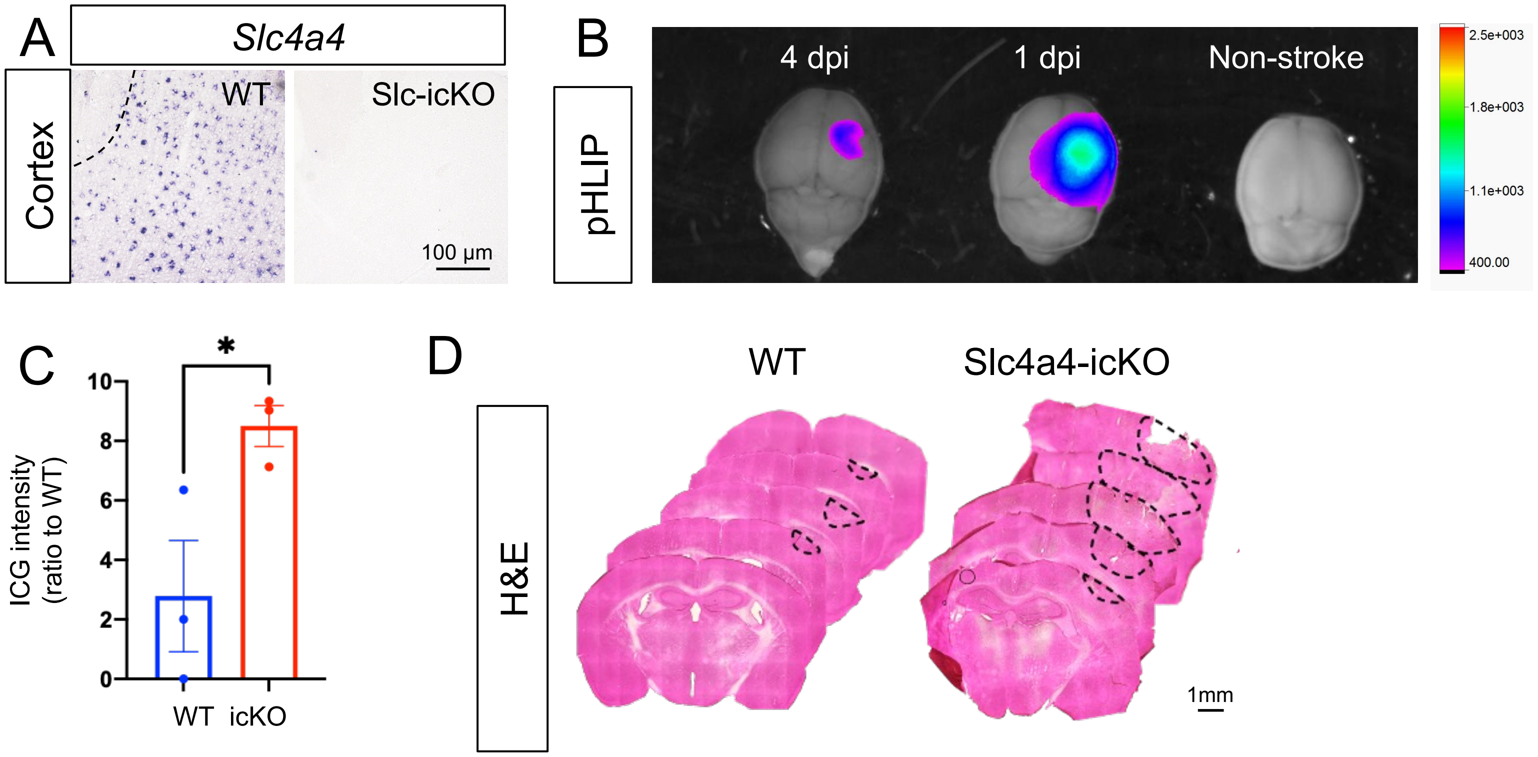


**Supplementary Figure 3. Slc4a4-icKO shows increased infarct size exacerbated acidosis and junctional loss after ischemic stroke. (A)** In situ hybridization confirms deletion of Slc4a4 in the cortex after stroke at 4 dpi. **(B)** Extracellular pH in the stroke lesions was measured by intraperitoneal injection pHLIP-ICG dye (1mg/kg). 24 hours after injection (1 and 4 dpi), brains from mice were harvested and imaged using the Bruker Xtreme Imager with 735 nm excitation and 830 nm emission wavelength. **(C)** Quantification of ICG intensity by fluorescence signal detected in WT and Slc4a4-icKO at 1 dpi. Each data point represents an individual animal. N = 3 animals per genotype. *p<0.05 by Student's t-test. **(D)** Representative images of serial brain sections (500µm apart) after H&E staining at 4 dpi.

**Supplemental Figure 4** Related to Figure 3

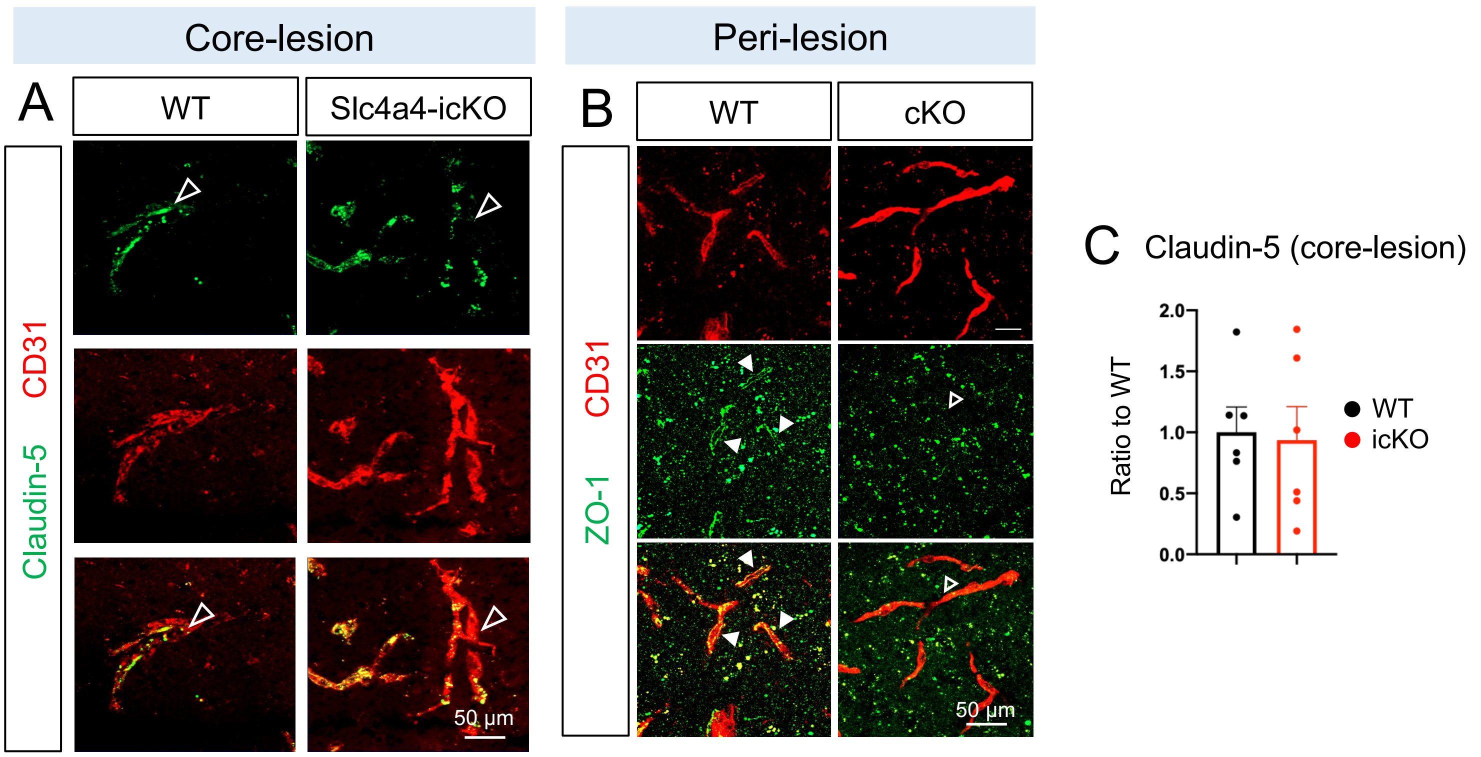


**Supplementary Figure 4. Slc4a4-icKO exacerbates BBB damage after ischemic stroke. (A)** Representative images of Claudin-5 colocalized with CD31 at the core-lesion area of WT and Slc4a4-icKO mice at 4 dpi. **(B)** Representative images of ZO-1 colocalized with CD31 at the peri-lesion area of WT and Slc4a4-icKO mice at 4 dpi. **(C)** Quantification of Claudin-5 colocalized with CD31 in the core-lesion. Data are presented as mean ± SEM. n = 6-8 blood vessels from N = 3-4 animals per genotype. ***p<0.001 by Student's t-test.

#### **Supplemental Figure 5** Related to Figure 4

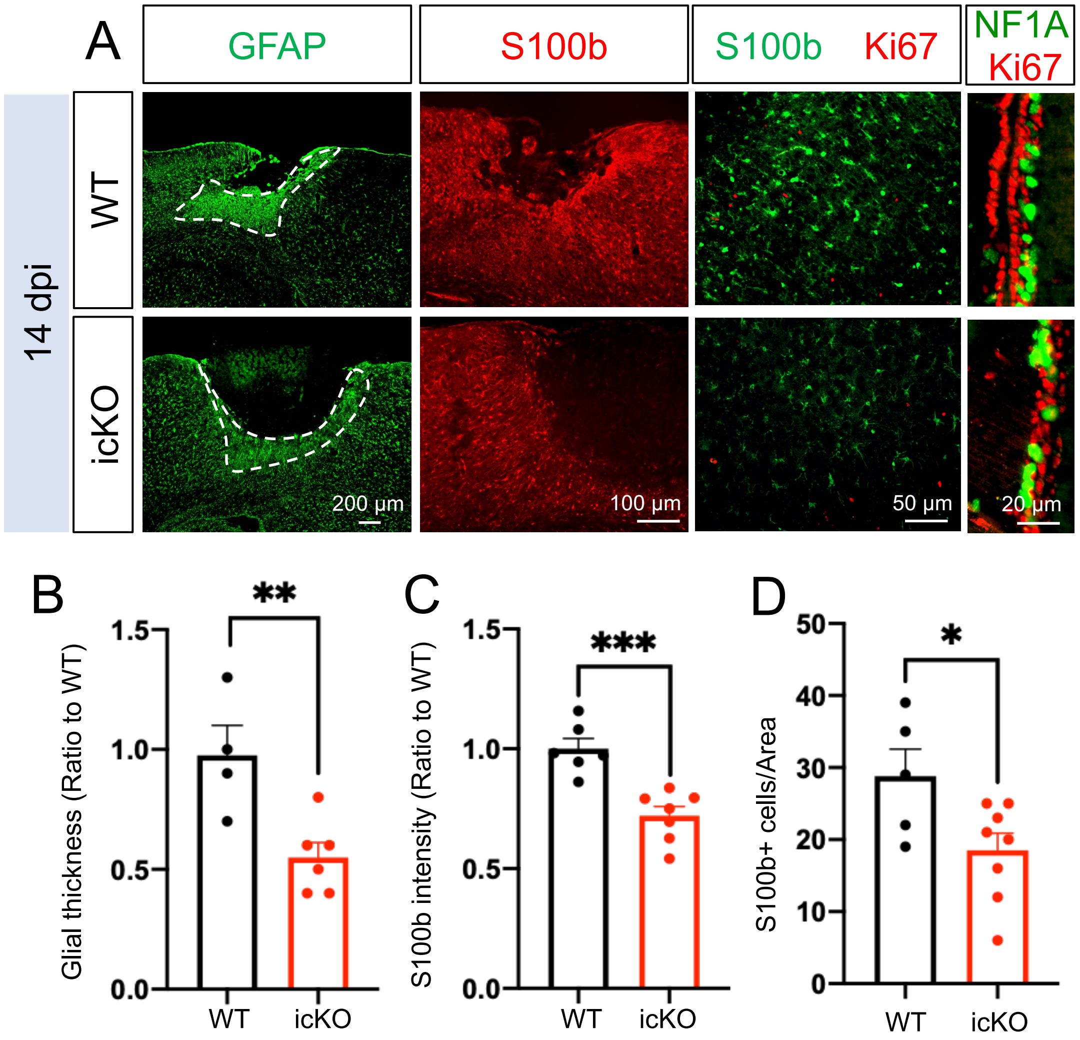


**Supplementary Figure 5. Loss of astrocytic Slc4a4 dampens reactive astrogliosis after stroke. (A)** Immunostaining of reactive astrocyte markers (GFAP, S100b) at the peri-lesion area and SVZ at 14 dpi. S100b+ cells are also co-labeled with proliferation marker Ki67 to indicate local astrocyte proliferation. **(B)** Quantification of glial thickness based on GFAP staining. **(C-D)** Quantification of S100b intensity and the number of reactive astrocytes based. Data are presented as mean ± SEM. Each dot represents an individual animal. N = 4-6 animals per genotype. *p<0.05, **p<0.01, ***p<0.001 by Student's t-test.

#### Supplemental Figure 6 Related to Figure 5

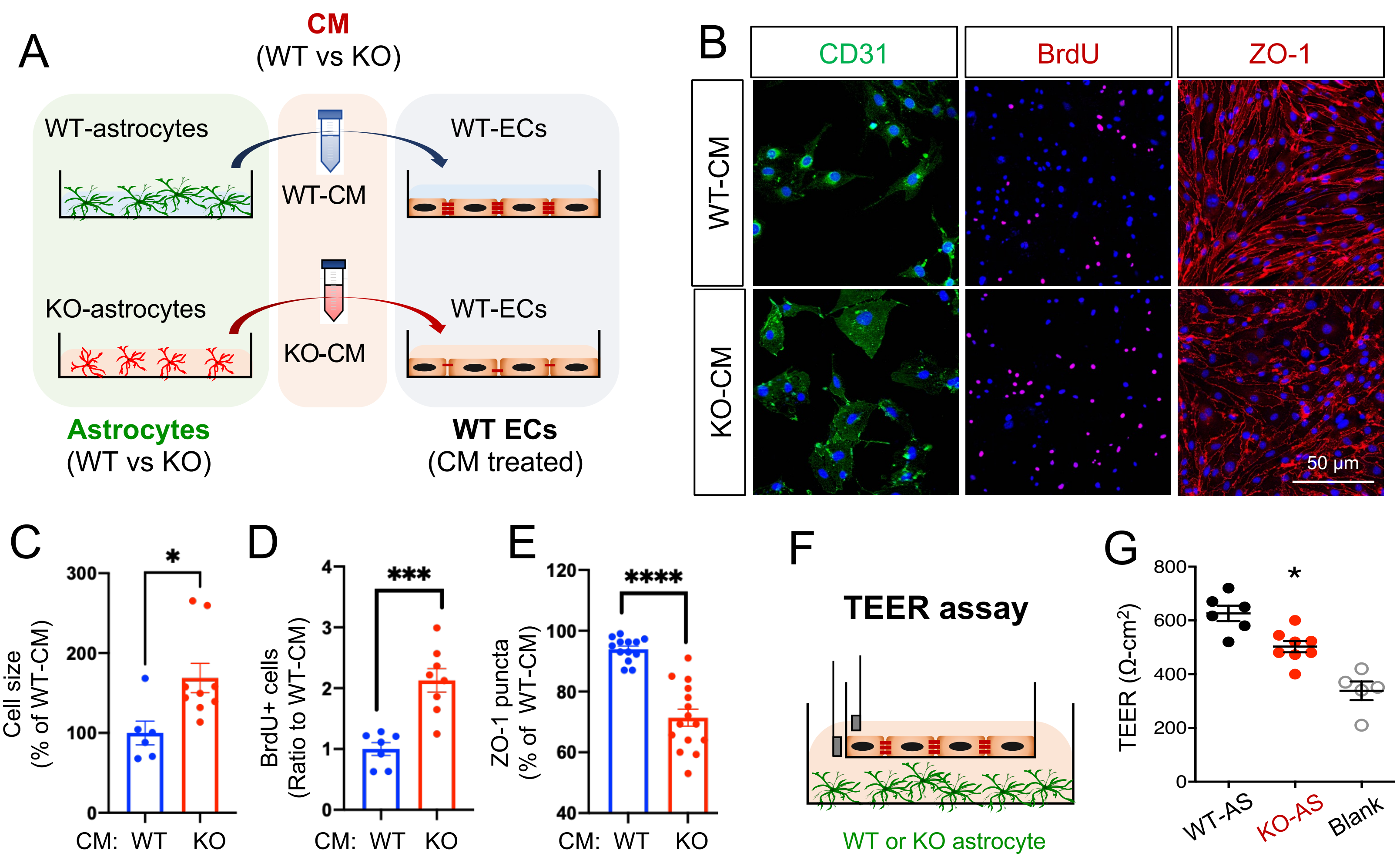


**Supplementary Figure 6. Slc4a4 regulates astrocyte-endothelia interaction via astrocyte derived cytokine CCL2 (A)** Mouse endothelial cell line (bEnd3) was incubated with conditioned media collected from primary WT and Slc4a4 KO astrocytes for 24 hours. **(B-E)** Endothelial cell size was examined by CD31 staining, cell proliferation was examined by BrdU assay, and tight junctional expression was examined by ZO-1 expression in bEnd3 incubated with conditioned media collected from primary WT and Slc4a4 KO astrocytes. Data are presented as mean ± SEM. Experiments were performed in duplicates for three independent assays. *p<0.05, ***p<0.001, ****p<0.0001 by Student's t-test. **(F)** Experimental setup of the transendothelial electrical resistance (TEER) assay. **(G)** The electro-resistance of the endothelial cell monolayer was measured as an indicator of the permeability of endothelial cells in the TEER assay. Data are presented as mean ± SEM. Wells without cells were used as blank control. Experiments were performed in duplicates for three independent assays. *p<0.05 by Student's t-test.

#### **Supplemental Figure 7** Related to Figure 5

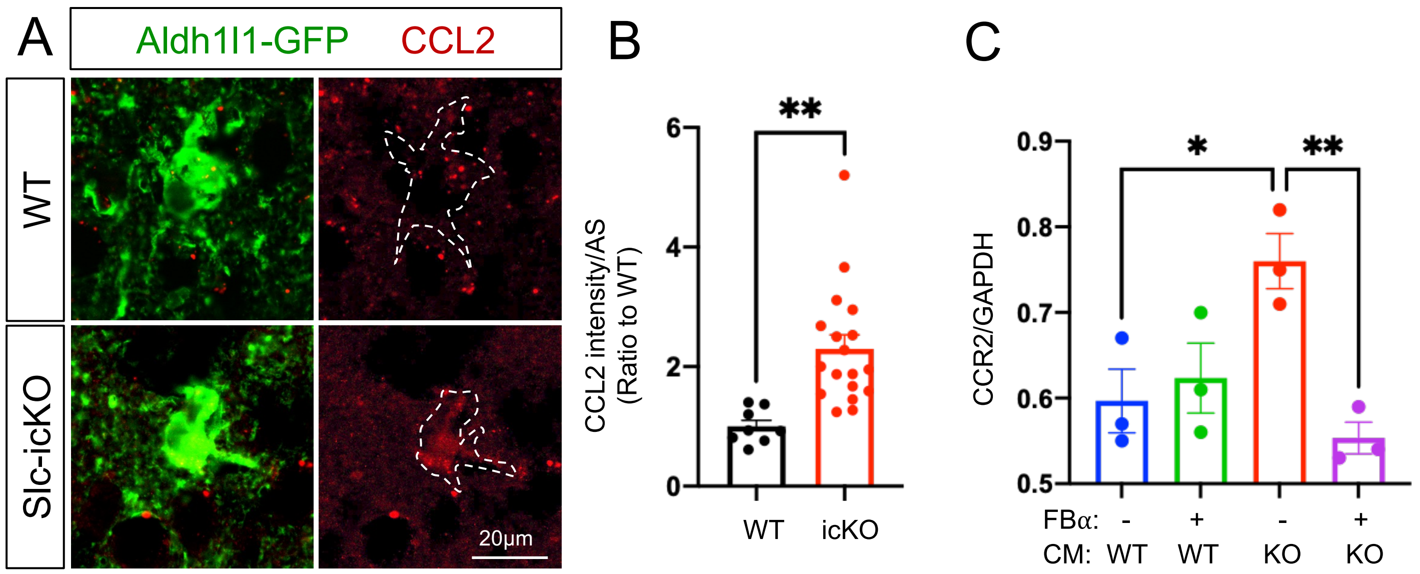


**Supplementary Figure 7. Loss of Slc4a4 upregulates astrocytic CCL2 and endothelial CCR2. (A-B)** Representative images and quantification of astrocytic CCL2 expression from double immunostaining of Aldh1l1-GFP and CCL2. Data are presented as mean ± SEM. n = 8-18 cells collected from N = 3-5 mice per genotype. **p<0.01 by Student's t-test. **(C)** Western blot quantification of CCR2 expression in bEnd3 cells incubated with WT- and Slc4a4 KO-CM with CCR2 FBα or control IgG. Data are presented as mean ± SEM. Each dot represents each independent culture. N = 3 independent assays. *p<0.05, **p<0.01 by two-way ANOVA.

**Supplemental Figure 8** Related to Figure 7

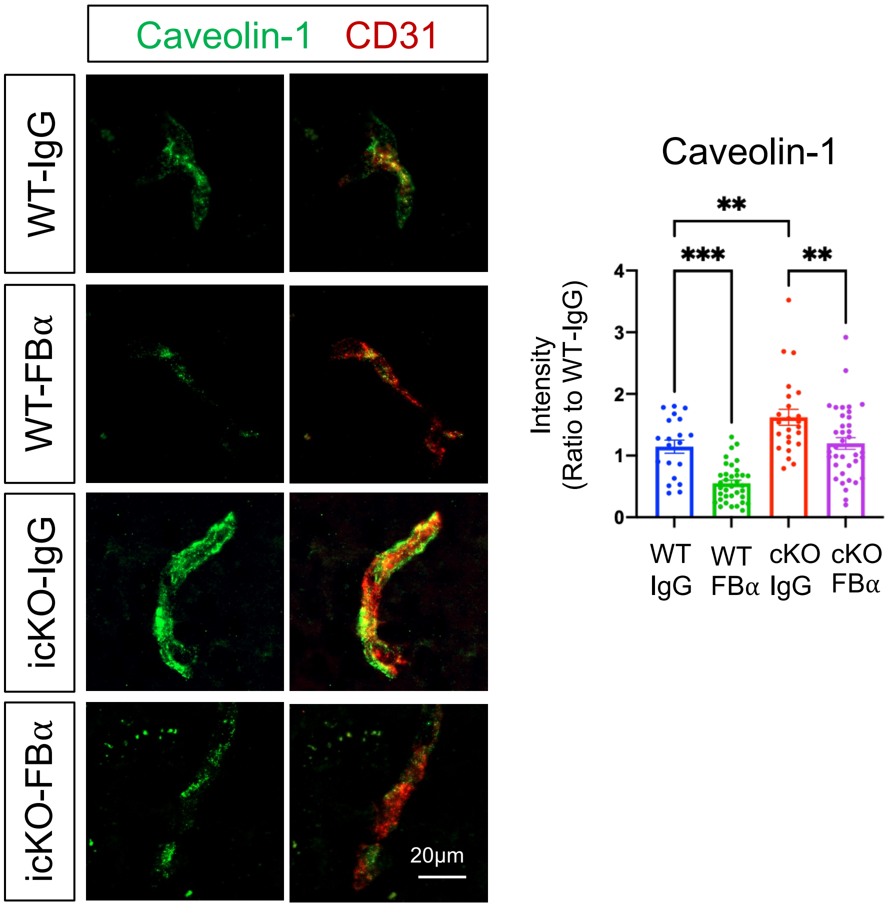


**Supplementary Figure 8. Loss of Slc4a4 upregulates endothelial caveolin-1.** Representative images and quantification of endothelial caveolin-1 expression from double immunostaining. Data are presented as mean ± SEM. n = 4-5 blood vessels per animal and N = 4-5 animals per group. **p<0.01, ***p<0.001 by two-way ANOVA.

#### **Supplemental Figure 9** Related to Figure 8

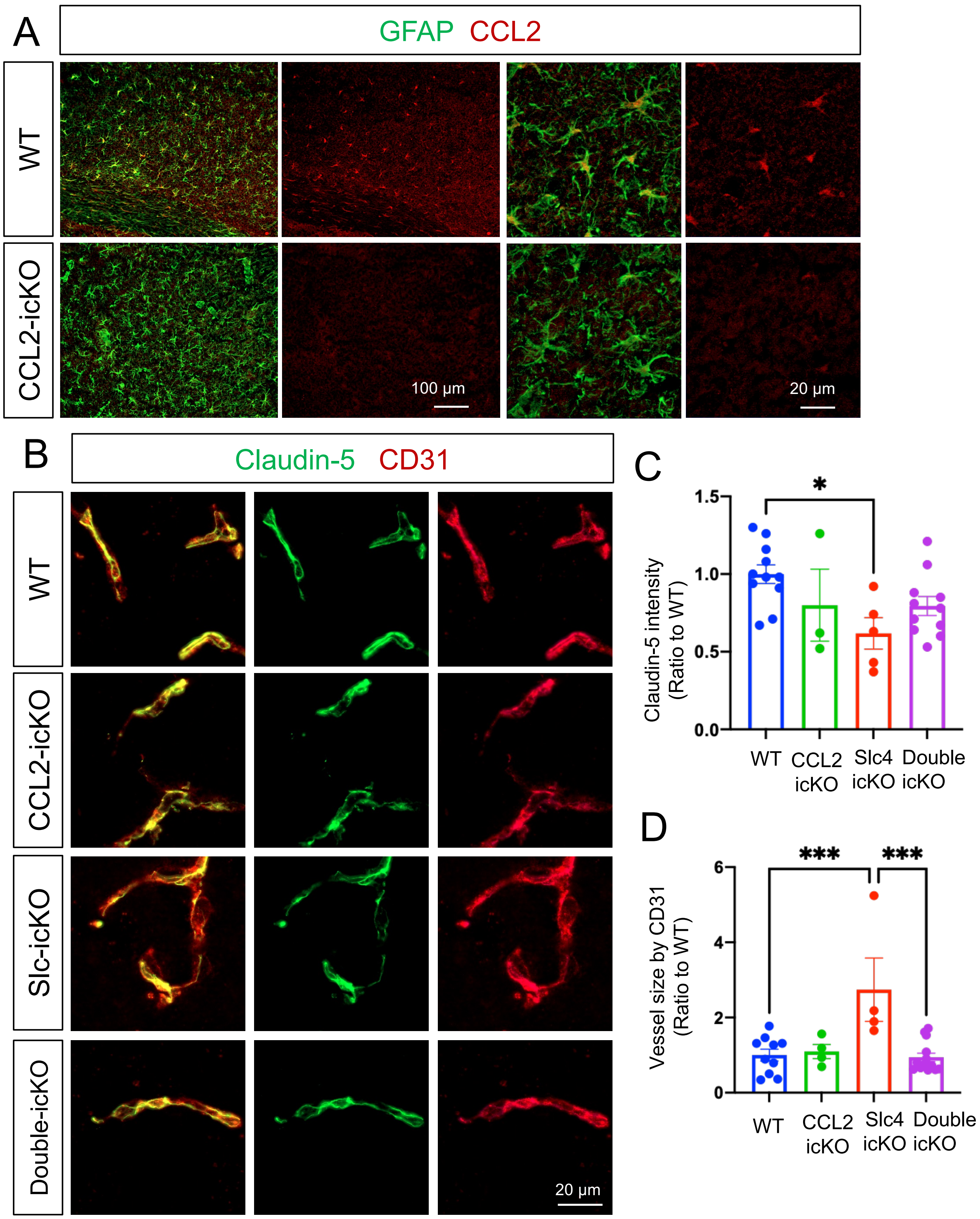


**Supplementary Figure 9. Deletion of astrocytic CCL2 rescues loss of Slc4a4-induced tight junctional marker expression. (A)** Confirmation of CCL2 deletion in reactive astrocytes in stroked CCL2-icKO mice by double immunostaining. GFAP was used as a reactive astrocyte marker. **(B)** Representative images of junctional marker expression (Claudin-5+; CD31+) in the cortex from WT, CCL2-icKO, Slc4a4-icKO, and double-icKO without injury. **(C-D)** Quantification of the intensity of Claudin-5 colocalized with CD31 and blood vessel size (CD31) from B. Data are presented as mean ± SEM. n = 1-2 blood vessels from N = 3-5 animals per group. $^{*}p<0.05$, $^{***}p<0.001$ by two-way ANOVA.

#### **Supplemental Figure 10** Related to Figure 8

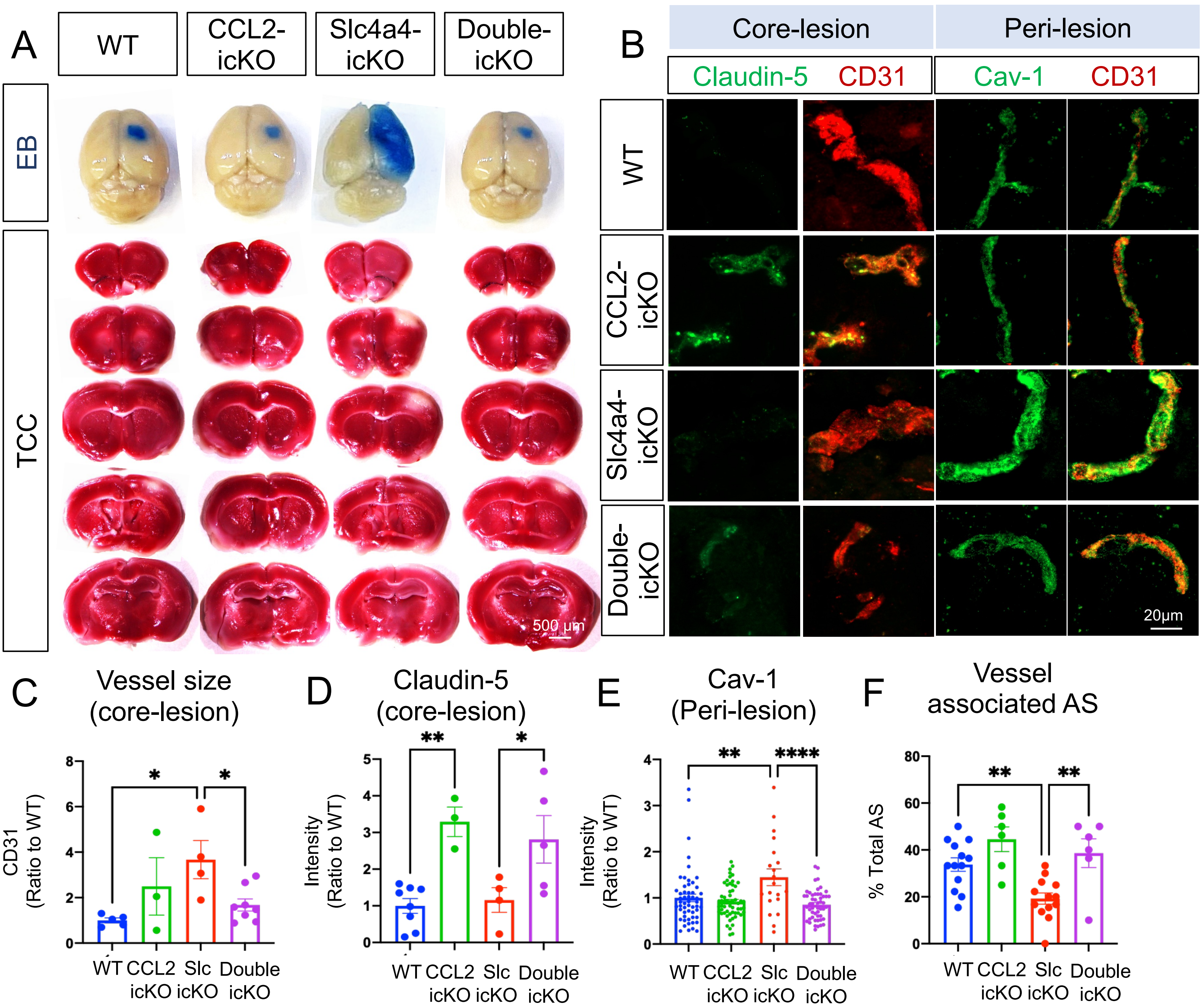


**Supplementary Figure 10. Deletion of astrocytic derived CCL2 rescues loss of Slc4a4-induced BBB damage. (A)** Representative images of Evans blue leakage and 2,3,5-Triphenyltetrazolium chloride (TTC) staining at 4 dpi. **(B)** Representative images of endothelial junctional marker (Claudin-5+; CD31+) and caveolae marker (Cav-1+; CD31+) at 4 dpi. **(C-D)** Quantification of blood vessel size and endothelial Claudin-5 expression from double immunostaining at the core-lesion by CD31. Data are presented as mean ± SEM. n = 1-2 blood vessels from N = 3-5 animals per group. *p<0.05, **p<0.01 by two-way ANOVA. **(E)** Quantification of the intensity of Cav-1 colocalized with CD31. Data are presented as mean ± SEM. n = 4-5 blood vessels from N = 4-6 animals per group. **p<0.01, ****p<0.0001 by two-way ANOVA. **(F)** Quantification of vessel-associated astrocytes by S100b and CD31 double immunostaining from main Figure 8E. Vessel-associated astrocytes are defined as those astrocytes whose somas occupy vascular territory. Data are presented as mean ± SEM. n = 4-5 blood vessels from N = 4-6 animals per group. **p<0.01 by two-way ANOVA.

#### **Supplemental Figure 11** Related to Figure 9

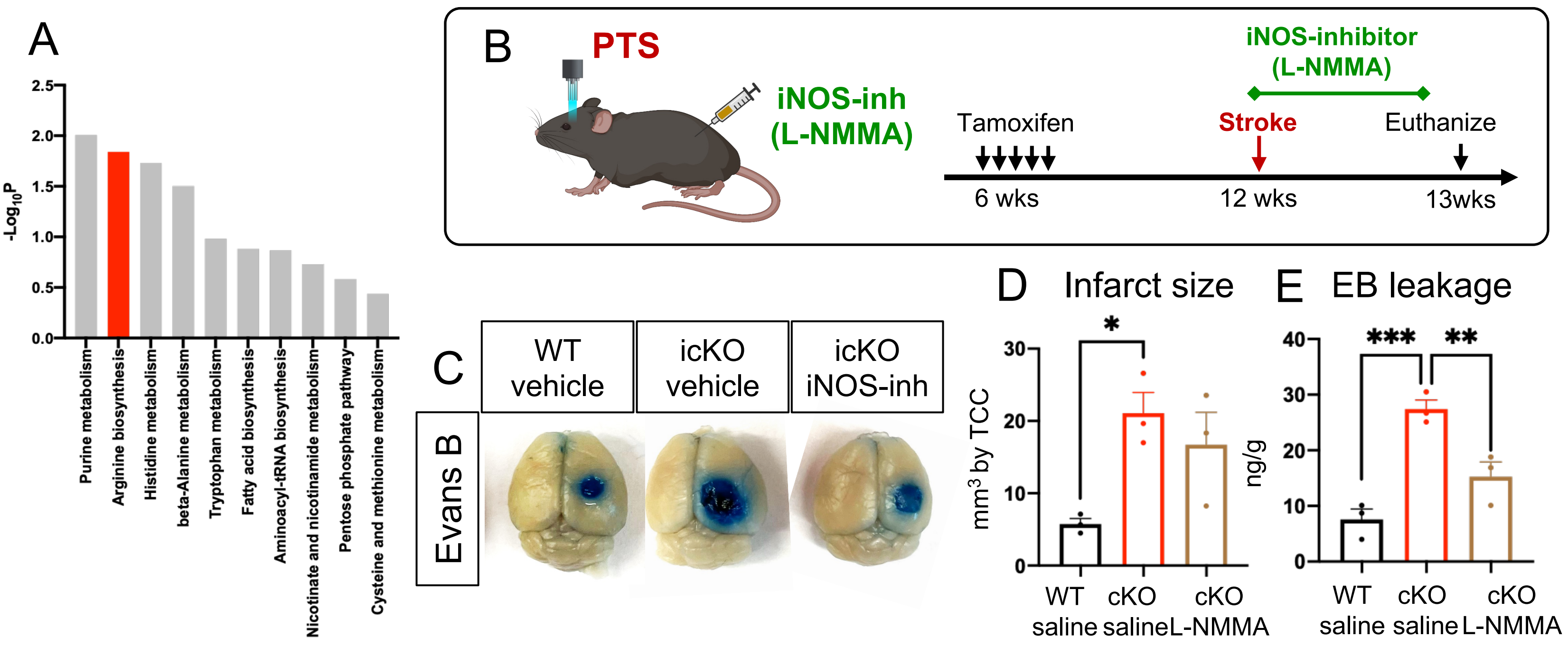


**Supplementary Figure 11. Inhibition of iNOS rescues exacerbated BBB leakage in Slc4a4-icKO after stroke (A)** Cortices was collected from WT and Slc4a4-icKO mice and subjected to unbiased metabolomic analysis, followed by pathway analysis. **(B)** Experimental scheme of the PTS induction in WT and Slc4a4-icKO mice, followed by daily intraperitoneal injection of iNOS inhibitor (L-NMMA) from 1-3 dpi. Brains were then harvested and analyzed at 4 dpi. **(C)** Representative images of albumin leakage (Evans blue) in the brain from WT and Slc4a4-icKO mice at 4 dpi with or without iNOS inhibitor treatment. **(D)** Quantification of infarct size based on 2,3,5-Triphenyltetrazolium chloride staining of serial 1mm-thick brain sections from stroked brains at 4 dpi. Data are presented as mean ± SEM. Each dot represents an individual animal. N = 3 animals per group. *p<0.05 by two-way ANOVA. **(E)** Quantification of Evans blue leakage by colorimetric assay from stroked brains at 4 dpi. Data are presented as mean ± SEM. Each dot represents an individual animal. N = 3 animals per group. **p<0.01, ***p<0.010 by two-way ANOVA
